# Steric Communication between Dynamic Components on DNA nanodevices

**DOI:** 10.1101/2022.12.15.520588

**Authors:** Y. Wang, S. Sensale, M. Pedrozo, C-M. Huang, M.G. Poirier, G. Arya, C.E. Castro

## Abstract

Biomolecular nanotechnology has helped emulate basic robotic capabilities such as defined motion, sensing, and actuation in synthetic nanoscale systems. DNA origami is an attractive approach for nanorobotics, as it enables creation of devices with complex geometry, programmed motion, rapid actuation, force application, and various kinds of sensing modalities. Advanced robotic functions like feedback control, autonomy, or programmed routines also require the ability to transmit signals among sub-components. Prior work in DNA nanotechnology has established approaches for signal transmission, for example through diffusing strands or structurally coupled motions. However, soluble communication is often slow and structural coupling of motions can limit the function of individual components, for example to respond to the environment. Here, we introduce a novel approach inspired by protein allostery to transmit signals between two distal dynamic components through steric interactions. These components undergo separate thermal fluctuations where certain conformations of one arm will sterically occlude conformations of the distal arm. We implement this approach in a DNA origami device consisting of two stiff arms each connected to a base platform via a flexible hinge joint. We demonstrate the ability for one arm to sterically regulate both the range of motion as well as the conformational state (latched or freely fluctuating) of the distal arm, results that are quantitatively captured by mesoscopic simulations using experimentally informed energy landscapes for hinge-angle fluctuations. We further demonstrate the ability to modulate signal transmission by mechanically tuning the range of thermal fluctuations and controlling the conformational states of the arms. Our results establish a communication mechanism well-suited to transmit signals between thermally fluctuating dynamic components and provide a path to transmitting signals where the input is a dynamic response to parameters like force or solution conditions.

## INTRODUCTION

DNA nanotechnology^1,2^ has garnered increasing interest for the development of nanoscale robotic systems due to the precise control over geometry afforded by this approach, the ability to design devices with complex motion^3^, and tunable mechanical properties^4^. In addition to these design features, advanced robotic devices and materials typically integrate functions like actuation, sensing, and communication, which combined can enable advanced capabilities like feedback control or autonomous behavior. While actuation^5–11^ and sensing^12–16^ have been widely demonstrated within the context of DNA nanotechnology^17^, communication – which entails transmitting information from one part of the system to another – has been much less studied. Many natural biomolecules have evolved mechanisms for signal transmission. One example of such mechanisms is allosteric communication within proteins, where local binding events regulate functions at distal locations through conformational changes or modulation of dynamic properties such as amplitudes of motion or conformational distributions^18–21^. Inspired by protein allostery, here we develop a mechanism for transmitting information via steric interactions between dynamic components within a DNA nanostructure.

Prior efforts have implemented approaches to transmit signals within DNA nanostructures either via transport of components, such as single strands^22–24^ or larger constructs^25–29^, along predefined tracks, or via conformational changes of the nanostructure resulting from structurally coupled motions^30,31^. The first of these approaches introduces hybridization and dissociation of nucleotide sequences as the driving mechanisms for communication^32^. These mechanisms enable very precise communication, due to the sequence specificity of DNA, at the expense of speeds and distance. Other studies have leveraged the precise geometric design of DNA origami^33,34^ to demonstrate larger motions in nanostructures exhibiting conformational changes involving stiff links coupled by flexible joints^7,8^ to achieve kinematically constrained motion, or in devices where conformational changes propagate among small repetitive structural units^30,35^. Such deployable structures allow for the transfer of forces and motions among large DNA components, which have been used to control molecular interactions^31,36^, and can be combined with triggering events, such as local enzymatic modifications or binding of an input molecule, to regulate device properties at distal locations^36,37^. Introduction of such conformational changes enables for allosteric communication. However, structurally coupling the motion of components removes degrees of freedom and limits the overall design flexibility.

In this work, rather than transferring motion through kinematic constraints or coupling local conformational changes, we establish an approach to transmit mechanical signals through the steric interaction among thermally fluctuating components. These interactions enable fluctuating components to regulate the dynamic properties of distal ones, such as amplitudes of motion or thermodynamics of binding interactions. Steric interactions have recently been demonstrated to constrain rotational motions of tight-fitting components in DNA origami rotary mechanisms^38^, including facilitating processive rotation in nanoscale motors^39^. Here we establish the ability to convey steric interactions between two distal thermally fluctuating components. We present a nanodevice comprised of two fluctuating arms connected to a base platform, where the arms are long enough to come into contact. We modulate the steric interactions by engineering the conformational distributions of individual arms and show that the conformational fluctuations of each arm can influence the dynamic motion of the other arm. In addition, we introduce binding interactions between arms and base platform to further control the conformational distribution of the individual arms and demonstrate that each fluctuating arm can also modulate the equilibrium binding distributions of the distal one. These results establish steric interactions as a mechanism for transmitting mechanical signals between thermally fluctuating components and lay a foundation for allosteric devices where local events regulate dynamic properties at distal locations.

## RESULTS AND DISCUSSION

### Design and Fabrication of a DNA Origami Device with Interacting Arms

A DNA origami nanodevice comprised of two fluctuating arms connected to a base platform via flexible hinges was designed in MagicDNA^40^. To allow for steric interactions, the arms were designed to be longer than half the length of the base platform, with lengths of left arm *l*_*L*_ = 41 nm, right arm *l*_*R*_= 46 nm and base platform arm *l*_*C*_ = 57 nm. The arms were designed with different lengths to study how length influences steric interactions. Each of the three components was made up of an 18-helix bundle organized into three layers of six helices. Two additional bundle layers were added to the bottom of the left-hand side of the base platform to allow for easy identification of the left and right arms in images (**Figure 1A**). The flexible hinge connection between both arms and the base platform comprises 6 ssDNA scaffold linkers. Three of these ssDNA connections were 2-nt-long pieces of scaffold (black strands in Figure 1A) arranged along a line defining the hinge axis of rotation, while the other three ssDNA connections were 30-nt-long DNA strands (blue strands in Figure 1A) introduced to facilitate tuning of the angular distribution of the individual arms^41^. At the ends of the bundles, poly-thymines (poly-T, either 5 or 10 T long) strands (orange strands in Figure 1A) were added to prevent base stacking of the arms with the base platform. We refer to this device as the “SteriDyn”, since it was designed to exhibit dynamic behavior regulated by steric interactions. In addition, the device contains sites for mutually complementary ssDNA overhangs on the arms and base platform to latch the left or right arms down to the platform (green and red strands). The design can harbor up to three latching overhangs for either arm to achieve high yield of latching (see **Figure 1B**). Structures were folded in 20 mM MgCl_2_ (gel electrophoresis shown in **Figure S3**), purified by PEG centrifugation^42^, and analyzed via TEM (**Figure 1B-C**).

**Figure 1.**
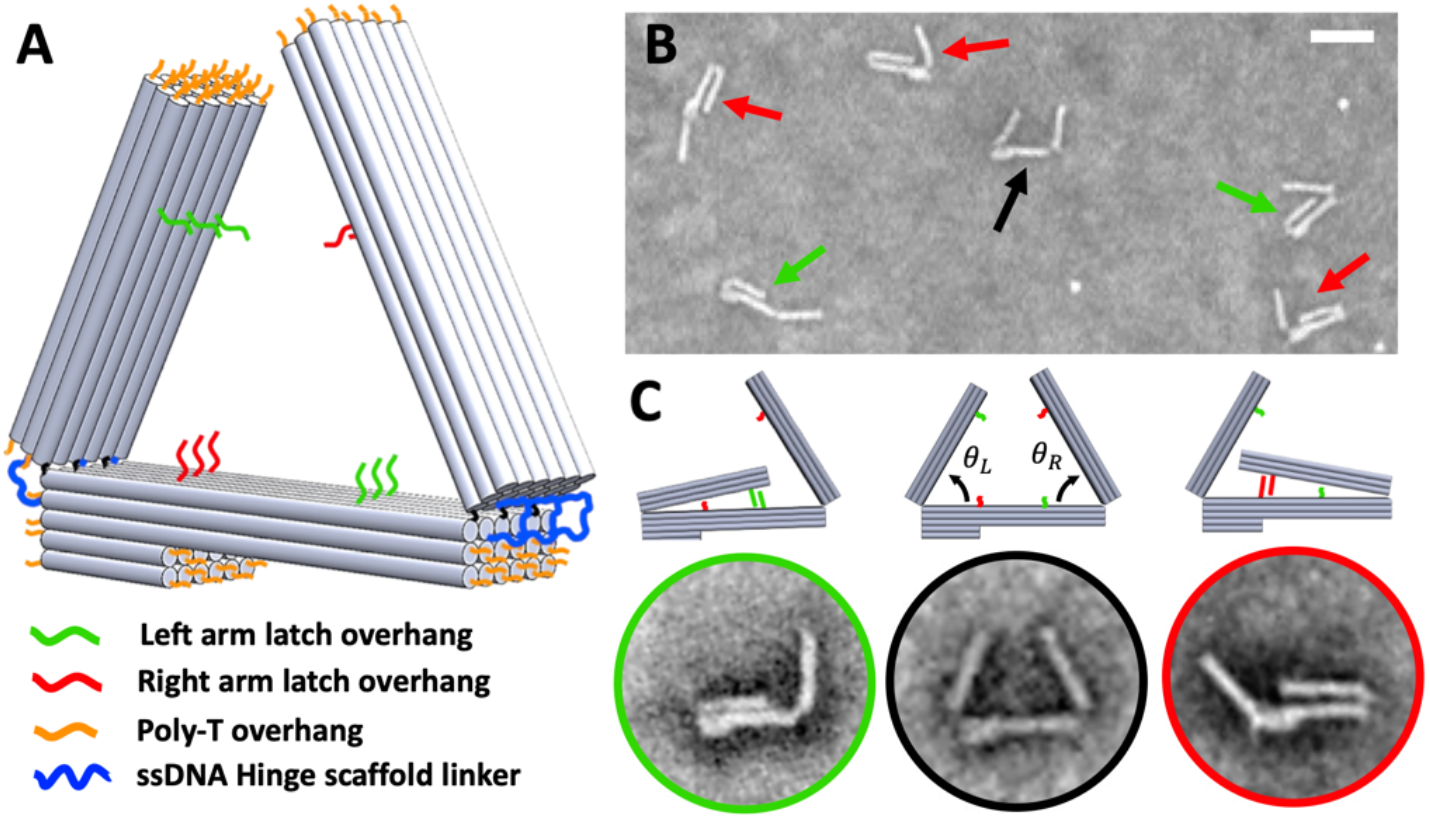
Design and fabrication of the DNA origami device. **(A)** Isometric view of our device with a 41-nm-long left arm, 46-nm right armB and 57-nm central arm. The central arm included an additional 6×2 square lattice structure at the bottom of one of its sides to distinguish the structure orientation. Green and red strands represent the extended overhangs introduced to latch the arms to the central arm. The hinge connection is made of 6 scaffold connections (3 black 2-nt-long ssDNAs and 3 blue 30-nt-long ssDNAs). Orange strands represent 5-nt-long poly-T overhangs. **(B)** TEM images of the devices show 3 typical conformations: left arm closed (red arrows), right arm closed (green arrows), and unlatched arms (black arrow). **(C)** Schematics and TEM images depict an example of the left arm latched by a green strut (green circle); an example of both arms open (black circle), with our convention for the left arm angle, *θ*_*L*_, and the right arm angle, *θ*_*R*_, shown; and an example of the right arm latched by a red strut (red circle). Scale bar: 50 nm.

### Hinge and Overhang Properties Control the Free Energy Landscape of the Arms

Prior studies have shown that the properties of the ssDNA linkers in DNA origami hinge joints modulate the flexibility of the hinge^7,41,43^. We first investigated SteriDyn devices with fluctuating arms where the hinge joint scaffold linkers were left single stranded and 5T overhangs were included on the ends of the arms and base-platform. We observed this baseline condition favors open conformations of the arms (see **Figures 2A** and **2F**). Under this “free hinge” condition, the arms behave independently since they both sample large angles where no steric interactions occur.

**Figure 2.**
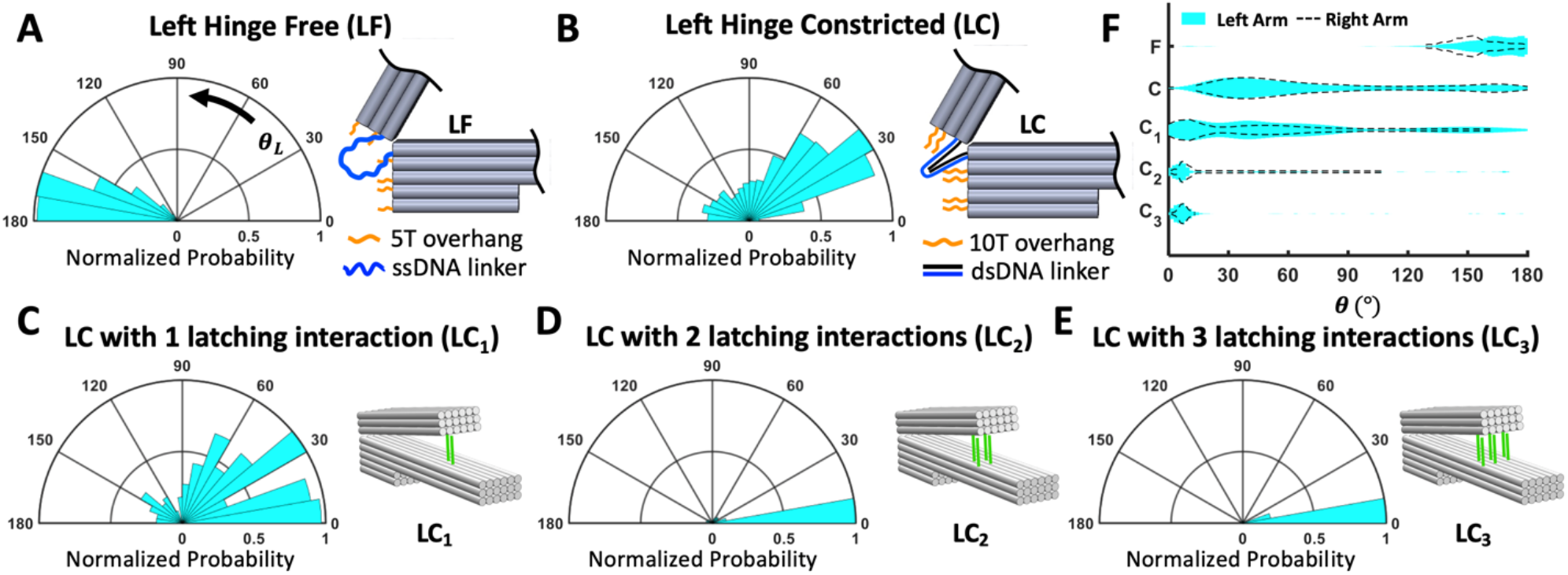
Engineering single arm angular distributions with scaffold connection and overhangs. **(A)** Baseline device with both left and right free arms (LF|RF, 160°|153° mean angles) results in open conformations. **(B)** Increasing the poly-T length of the left arm connections from 5 nt to 10 nt and base pairing these connections with complementary staples shifts the arm’s angular distribution towards more closed conformations (LC|RF, 74°|150°). **(C)** Modifying LC|RF by introducing the duplex DNA overhangs that latch the left arm shifts the distribution towards even smaller angles. LC_1_|RF (left), **(D)** LC_2_|RF (middle), and **(E)** LC_3_|RF (right) represent arms latched by 1, 2, and 3 pair(s) of overhangs, respectively. Angular histograms were normalized by setting a maximum bin height of 1. **(F)** Summary of the hinge angles, *θ*, for the left (cyan) and right (dashed black line) arms compared for all five cases where the opposing arm is in the free configuration (individual distributions for right arm shown in Figure S6). Outliers where p<0.02, where p is the normalized probability, were removed from the distribution for clear comparison (full distributions are shown in the Figure S7). Sample sizes N for LF|RF, LC|RF, LC_1_|RF, LC_2_|RF, and LC_3_|RF are 303, 521, 240, 193, and 180, respectively.

We then changed the properties of each arm individually, while leaving the other arm free to measure individual hinge properties. We first used a hinge design approach inspired by prior work^41^, where scaffold linkers were made double-stranded in two sections by extending 6 staples (3 from the arm and 3 from the base-platform) by 15 nt to base-pair the scaffold linkers. We also extended the poly-T overhangs from 5T to 10T to inhibit base-stacking interactions more strongly. This “constricted hinge” condition allowed us to shift the angular distributions of each arm towards smaller angles, and thereby more closed conformations, with a mean angle of 74° (**Figures 2B and 2F**). In addition to these dynamic conformations where arms can sample a range of angles, we designed sites both in the arms and platforms which allow for the placement of up to three overhangs capable of latching the arms onto the base platform. A single 8 bp latching overhang leads to a mixture of latched and unlatched states (**Figures 2C**), while two and three latching overhangs leads to nearly all arms exhibiting a latched conformation (**Figure 2D and 2E**). We ran similar experiments to demonstrate control over conformations of the right arm with the left arm being in the free configuration (**Figure S6**). **Figure 2F** shows a summary of the left and right arm conformations in each condition tested in the absence of steric interactions (i.e., other arm at open angles). The two arms exhibit similar mechanical properties for the designs tested.

### Steric Interactions Modify Energy Landscapes of Distal Dynamic Components

Leveraging these approaches to engineer individual arm conformations, we next attempted to engineer steric interactions between the arms. Geometrically, steric interactions should yield a region of the 2D angular conformational space where both arms cannot coexist due to steric clash (i.e., a “dead zone” in the conformational space). When both arms can *independently* adopt angles inside this conformational space, each arm will exclude the other. This steric competition should then modify the angular distributions sampled by the individual arms. We hypothesized that the behavior of SteriDyn devices with interacting arms could be predicted from individual arm properties (**Figure 1**) and steric interactions. While coarse-grained models such as oxDNA can reproduce the mechanical properties and predominant motions of DNA origami structures^43–46^, they cannot access the timescales required to obtain large data sets of possible conformations of complex DNA origami nanostructures, such as the SteriDyn. To this end, we constructed a minimalistic mesoscopic model of our structures that accurately captures the geometric features and motions involved in our system (**Figure 3A**).

**Figure 3.**
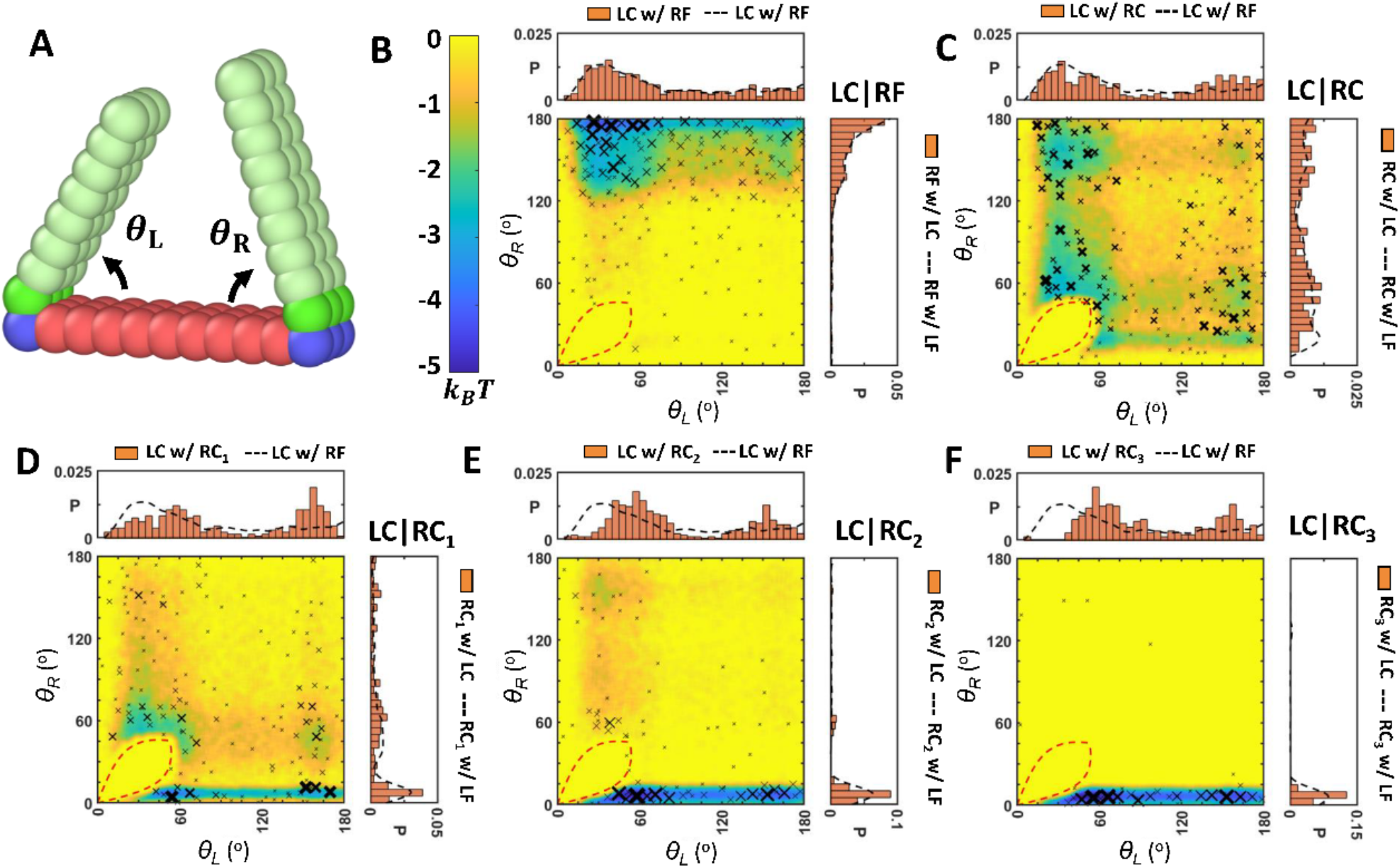
Angular conformational space of the interacting arms. **(A)** Schematic of our coarse-grained model and angle convention. The central arm (pink beads) of our device is functionalized to the two mobile arms (cyan beads) by means of physical hinges (involving blue and green beads as well as the closest arm beads) as shown in **Figure S8. (B-F)** Experimental counts (crosses) and simulated energy landscapes (color) of left (x axis) and right (y axis) angles for different systems of interacting arms. The area inside the dashed line represents the boundary of the “dead zone”. Experimental counts were clustered by means of a distance cutoff (7.5°) and the size of the crosses are logarithmically associated to the number of points in each cluster. Clusters were built from largest to smallest sizes to maximize the number of points in each cluster. Energy landscapes were constructed by Boltzmann inversion from 15 simulations, where measurements were taken every 200,000 steps on a total of 100 million steps, and smoothed by means of a mean filter over a rectangle of size 5°-by-5°. A single visit was associated with angular conformations not sampled in our simulations. Experimental results are also shown as histograms in marginal plots. Dashed lines are the reference for hinges experiencing no steric effects (open opposite arms). Sample size in experiment from B to F are 521, 383, 264, 356, and 232, respectively.

Briefly, our mesoscopic model treats the interacting arms as rigid bodies tethered to a rigid platform and capable of rotating around the hinge attachment points. The arms and the base platforms consist of spherical beads with a diameter of 6 nm (∼ 3 DNA helices) placed to resemble the geometry of the SteriDyn (a base platform with a length of 9 beads, left arm of length 7 beads and right arm of length 8 beads, with all three elements having a thickness of 1 bead and a width of 3 beads). The two arms are capable of rotating around their axes by means of experimentally informed energetic penalties *U*(*θ*) applied to the corresponding hinges. These penalties, which implicitly model the effects of hinge designs and latching overhangs, were chosen to follow the Boltzmann distribution *U*(*θ*) = −*k*_*B*_*T* ln *P*(*θ*) + *U*_0_, where *P*(*θ*) is the experimentally measured probability density associated with finding a chosen arm at a particular angle *θ* (in the absence of interactions with the other arm), and *U*_0_ is a shift factor that makes *U*(*θ*) = 0 at the angle of minimal energy (or maximum probability density). Steric interactions between the interacting arms and between these arms and the rigid platform were introduced by means of short-range repulsive potentials, which allowed us to reproduce the steric exclusion between the mobile components of our devices.

We first tested this model for a design where the left arm hinge is constricted (LC) and the right arm hinge is free (RF); note that this case is the same as that presented in **Figure 2A** and represents a structure with negligible steric interactions. Langevin dynamics simulations of this structure were performed, and the free energy landscapes were reconstructed from angular distributions sampled by the two arms using Boltzmann inversion from 15 independent simulations, where measurements were taken every 200,000 time steps for a total of 100 million steps. The color contour map plotted in **Figure 3B** depicts the 2D angular free energy landscape constructed from simulations. To compare these predictions against experiments, we overlayed onto this map the set of angles sampled by our SteriDyn device as measured from the TEM images. These angles are indicated by black crosses (X), where the size of the crosses indicates the number of events at a particular location. Experimental counts were clustered by means of a distance cutoff (7.5°) and the size of the crosses are logarithmically associated to the number of points in each cluster. The experimentally observed conformations of the device arms are consistent with the predicted energy landscape (i.e., most events fall in the low energy regions). 1D histograms depicting the experimental angular probability distributions of each arm (**Figure 3B**, marginal plots) also agree well with the corresponding 1D angular distributions obtained from simulations in absence of steric interactions (**Figure S10**).

To engineer steric interactions between the two dynamic arms, we constructed SteriDyn devices with both left and right hinges in the constricted condition (LC and RC, respectively), which cause the arms to adopt relatively small angles. We again observe that the experimentally measured conformations are in good agreement with the simulation predictions (**Figure 3C**, and **Figure S10**). Both arms exhibit a wide range of angles including small angles, but the conformational distribution reveal the clear presence of a sterically clashed “dead zone”, which can be defined by the geometry of the two arms (red dashed line, see Methods section for detailed calculation). The dead zone is asymmetric due to the difference in length between the arms, showing that the longer right arm indeed excludes more conformational space of the shorter left arm than *vice versa*. Focusing on the left arm, the depletion of small angle configurations due to the sterically clashed dead zone results in an increase in large angle conformations, leading to a bimodal distribution with a small angle population and a large angle population observed in the 1D *θ*_*L*_ distribution. The 2D angular distribution reveals that the large left-arm angles (*θ*_*L*_) corresponds primarily to small right-arm angles (*θ*_*R*_) less than ∼60° (angles where right arm can interact with the left arm); whereas for *θ*_*R*_ larger than ∼60°, we observe the majority of *θ*_*L*_ at small angles. The 1D angle distribution of the left arm (**Figure 3C**, top) similarly reveals a depletion of small-angle configurations and a corresponding increase in large-angle configurations.

Based on these results, we reasoned that forcing the right arm to be at small angles would yield stronger steric interactions to alter the left-arm angle distribution. Hence, we constructed various SteriDyn devices where the left arm was constricted (LC) and the right arm was constricted (RC) and could further latch to the base-platform via 1, 2, or 3 overhang attachments, termed RC_1_ (**Figure 3D**), RC_2_ (**Figure 3E**), and RC_3_ (**Figure 3F**), respectively. We again observe a clear sterically clashed dead zone and a stronger shift towards larger angles of the left arm, *θ*_*L*_, (with corresponding depletion of small angles). Interestingly, LC|RC1, where the right arm exhibits both latched and unlatched states, results in the largest increase in the population of devices with *θL* larger than 120°. LC|RC2 and LC|RC3, which are nearly all latched, cause a clear depletion of angles below ∼40°, but when the right arm is latched (i.e., *θR* ≤ 15°) the left arm can rotate down on top of the right arm, leading to several devices that accumulate near the lower envelope of the steric dead zone. These results demonstrate the capacity to transmit a signal (i.e., influence conformational distributions) between distal dynamic components *via* steric interactions.

### Steric Interactions Modulate Binding States of Distal Dynamic Components

Besides concerted conformational shifts, allosteric mechanisms can also control binding/unbinding reactions in distal components^21,47–49^. Here we studied how a range of SteriDyn arm constraints could regulate the binding state (i.e., latched vs. unlatched) of the opposing arm. Focusing on the left arm, we constructed SteriDyn designs with a wide range of latching efficiencies (**Figure 4**). We defined an arm as “latched” if it adopts a configuration of *θ* ≤ 15°. Hence, even with no overhangs, it is possible for an arm to appear latched, although with low probability (<3%). We tested designs where the left arm is constricted without latching connections (LC), with one or two latching connections (LC_1_ and LC_2_), and two versions of the LC with three connections: LC_3_ introduced earlier and one with a higher fraction of GC base-pairs (LC_3(HGC)_) in the latching connections. We used the constricted right arm (RC) as the baseline condition. These designs ranged from 2% to 97% in the fraction of latched left arms (**Figure 4**).

**Figure 4.**
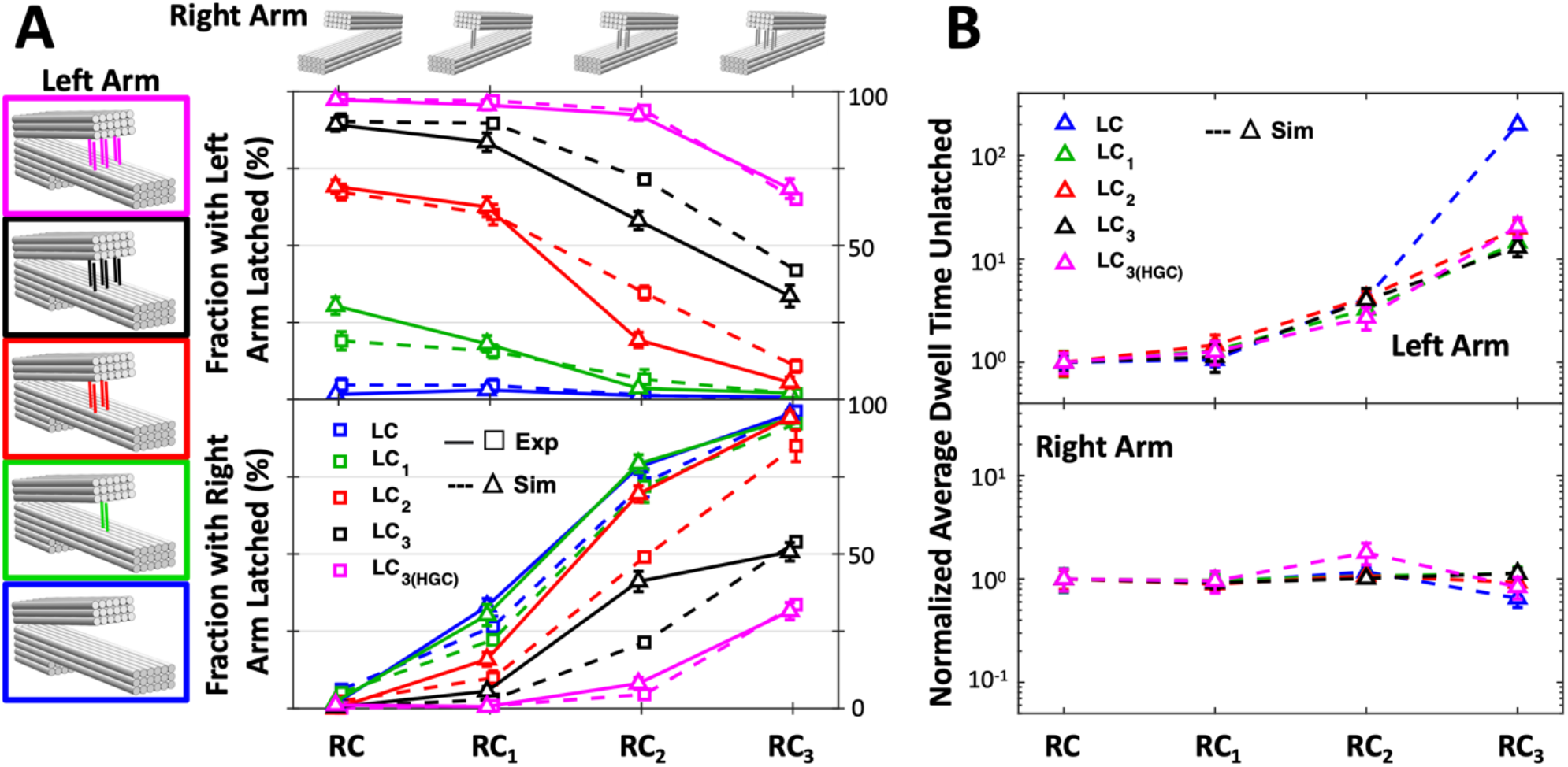
A) Simulated (dashed - squares) and experimental (solid - triangles) percentages of left (top plot) and right (bottom bot) latched arms for different numbers of overhangs. Simulation and experimental data were slightly offset from each other in the horizontal direction for visibility. A threshold θ=15_°_ was considered for the latched state. The x-axis is associated with right arm designs RC, RC_1_, RC_2_ and RC_3_, from lower to increasing propensity for latching. The error bars for experimental data were calculated by bootstrapping (see methods for details). The error bars for simulation data were calculated from the standard error associated to 15 simulations of each device (50 for strongly interacting arms). Error bars that are smaller than the symbols are not shown. B) Normalized dwell times in the unlatched state for the left arm (top plot) and right arm (bottom plot) obtained from simulations, where the average dwell time for each design was normalized by the case with no latching of the right arm (i.e., all cases were divided by LC_X_ paired with RC), showing the increased latching affinity of the right arm led to an increase in unlatched dwell times of the left arm, but does not affect unlatched dwell times of the right arm. The error bars were calculated from the standard error associated to 15 simulations of each device (50 for strongly interacting arms).

To regulate the binding state of the left arm through steric interactions, we constructed designs where the right arm could also latch to the bottom platform via one, two, or three interactions (RC_1_, RC_2_, and RC_3_). We find that due to steric interactions, latching of the right arm prohibits latching of the left arm. For each left arm design (LC, LC_1_, LC_2_, LC_3_, and LC_3(HGC)_), the propensity of left arm latching decreases with increasing latching affinity of the right arm. Low affinity latching of the right arm (RC_1_) causes a minor decrease in latching across all left arm designs. However, adding two latching overhangs to the right arm (RC_2_) leads to sharper drops in the left arm latching propensities for all designs except LC_3(HGC)_. Finally, adding three latching overhangs on the right arm (RC_3_) leads to continued decreases in the left-arm latching of LC, LC_1_, and LC_2_ and a significant drop in the left arm latching of LC_3(HGC)_. These results show that dynamic arm components can regulate the binding interactions of distal components through geometric design that mutually excludes binding states.

Interestingly, we observed cases where despite both arms latching less than 50% of the time, we still observed steric communication between the latching arms. For example, RC_1_ decreased the latching of LC_1_ from 30% (when paired with RC) to 18% (when paired with RC_1_) even though RC_1_ individually (i.e., when paired with LC) only exhibited 32% latching. We hypothesized that this steric communication was due to modulation of binding kinetics; that is, left arm latching sites being occluded part of the time due to binding of the right arm would decrease the latching rate of the left arm. To explore this kinetic modulation, we modified our Langevin dynamics simulations by introducing experimentally informed binding and unbinding probabilities *p*_on_ and *p*_off_ between the arms and the platform at an angle *θ*_0_, so that the dynamics for *θ* > *θ*_0_ (“open state”) follow the energy landscape *U*(*θ*) of the corresponding arm in absence of overhangs, while the dynamics for *θ* ≤ *θ*_0_ (“latched state”) are consolidated into a partially absorbing boundary. We then quantified dwell times in the unlatched and latched states from these simulations. As we do not have experimental data on absolute dwell times, we report relative dwell times, normalizing to the baseline case where there is no latching of the other arm. For example, we calculated the average dwell times from simulations in the unlatched state for each LC_1_ design and divided by the average dwell time for the LC_1_ paired with RC (i.e., no latching of the right arm). These results illustrate that increasing latching affinity of the right arm increases dwell times of the left arm in the unlatched state (**Figure 4B**) and does not influence the dwell times of the left arm in the latched state (**Figure S12**). Thus, dynamic binding of structural components can sterically modulate the binding kinetics of distal dynamic components.

Our results also show that for cases with similar latching affinity of the left and right arms (e.g., LC_1_|RC_1_, LC_2_|RC_2_, and LC_3_|RC_3_), the right arm outcompeted the left arm to achieve a higher latching efficiency. For example, for the SteriDyn device combining LC_1_|RC_1_, the right arm latched with 32% efficiency and the left arm latched with 18% efficiency. This asymmetry results from minor differences in the flexibility of the left and right hinges, as observed in the free-arm angle distributions (**Figure 2C**) and in the binding kinetics of individual arms, as the right arm requires a lower binding constant to be found in its closed conformation as often as its left counterpart (**Table S2**).

## CONCLUSIONS

We demonstrated a mechanism to transmit information between distal dynamic components of DNA nanodevices via steric interactions. In particular, we designed a dynamic device with three components, a base platform and two dynamic arms, whose geometries were chosen to allow for steric interaction between the opposing arms. We also exploited recently established approaches to modulate the flexibility of hinge joints to design either hinges that stayed mostly open, to avoid interactions between the arms, or hinges that stayed mostly closed, to promote steric interactions. Finally, we added overhangs between the arms and base platform to provide binding sites for dynamic conformational changes between latched and unlatched states.

We demonstrated the utility of steric interactions for the modulation of conformational landscapes of distal components both by influencing range of motion and distributions of binding states. Our mesoscopic simulations revealed that steric modulation can be accurately predicted from individual arm properties, and the experimentally validated simulations also allowed us to explore the full 2D conformational (and free energy) landscapes. We found that the interactions between the arms depended on specific structural, mechanical, and dynamic design parameters. For example, the larger length of the right arm led to an asymmetry in excluded angles as apparent in the 2D conformational distributions, and the balance of competing latching interactions depended on the number and sequence of the latching overhangs. Our simulations further revealed that the steric interactions lead to kinetic modulation of binding states through occlusion of binding sites.

These results establish steric interactions as a useful approach to transmit information (e.g., motion or binding) between distal dynamic components. The fact that these components remain dynamic suggest they can still be sensitive to local environmental inputs such as ion concentrations^50^, temperature^51^, or forces^52^. Hence, similar designs could convert environmental inputs into steric signals that regulate function. The possibility of designing systems with more than two dynamic arms, or even arrays of dynamic arms, could provide a foundation for leveraging steric interactions to perform logic steps^53^ or transmit signals over longer distances. While such tasks have previously been accomplished by means of strand displacement reactions, steric interactions promise robust and faster dynamics over larger length scales and environment sensitivity.

## METHODS

### Fabrication of the DNA Origami SteriDyn

The SteriDyn device was designed using the MagicDNA software^40^. An 8064 nt single-stranded scaffold was employed (M13MP18 bacteriophage virus prepared in our laboratory as previously described^54^. The staple sequences were generated and subdivided based on the modules of the device (e.g., left arm, central arm, right arm, hinge vertex, and latching struts). The process of design in MagicDNA and structure connectivity overview are shown in **Figure S1-S2**. The staples were ordered from a commercial vendor (IDT, Coralville, IA, **Tables S3-S4**). Prior to folding, staples were selected based on the desired modifications.

Folding reactions were carried out with 20 nM scaffold and 200 nM of each staple strand in a ddH2O solution containing 5 mM Tris, 5mM NaCl, 1 mM EDTA, and 20 mM MgCl2, at pH 8.0. An initial test of MgCl2 concentrations revealed 20 mM yielded quality folding (**Figure S2**). These reactions were subjected to thermal annealing in a thermal cycler (Bio-Rad, Hercules, CA) consisting of rapidly heating the solution to 65°C for 10 min, followed by annealing from 61°C and 46°C for 3 hours per degree Celsius, from 45°C and 24°C for 30min per degree Celsius and then cooling for 30 min at 4°C.

### Purification of SteriDyn

The structures were purified by centrifugation in a polyethylene glycol (PEG) solution^42^. PEG purification was performed by mixing 50 uL PEG buffer (15% PEG MW8000, 200 mM NaCl, and 100 mM Tris) with an equal volume of folded DNA origami structures followed by 16,000 g centrifugation for 30 min. Structures were then resuspended in 0.5xTBE with net 10mM MgCl2 to a concentration of 20 nM as quantified by measuring UV absorption on a NanoDrop (NanoDrop 2000C Spectrophotometer, Thermo Scientific). The structures were diluted to 1 nM (using the same buffer, 0.5xTBE with net 10mM MgCl2) for TEM imaging for a reasonable particle density and avoiding interactions among structures.

### TEM imaging and experimental data analysis

For TEM grid preparation, 4 μL of sample volume was deposited on Formvar-coated copper TEM grids, stabilized with evaporated carbon film (Ted Pella; Redding, CA). The sample was incubated on the grid for 4 min and then wicked away with filter paper. The sample was then stained by applying 10 μL 2% uranyl formate (SPI, West Chester, PA) twice for 1 second and 5 seconds, respectively, and the stain solution was wicked away after each incubation with filter paper. TEM imaging was carried out at the OSU Campus Microscopy and Imaging Facility on an FEI Tecnai G2 Spirit TEM at an acceleration voltage of 80 kV at a magnification of 45,000x. The raw TEM images (**Figure S4**) were first organized into galleries (**Figure S5**) using the EMAN2 software (v2.3) particle picking feature, which streamlined the angle measurement process. Approximately 200 particles were measured for each SteriDyn condition for lowering uncertainty (**Table S1**). Angles of each particle were measured manually using the software ImageJ by picking four points (1 at left inner arm end, 1 at left hinge vertex, 1 at right hinge vertex, and 1 at right hinge inner end) directly on the particle. The hinge angles can be simply solved by trigonometry. We used MATLAB as the post-processing tool to convert the angle data sets to probability density histograms. For getting the error bar of SteriDyn latching fraction, we bootstrapped raw angle distribution data 50 times (sampling with replacement). The fraction number was calculated for every bootstrapped data (50 fractions) and its standard deviation was used as error bar^55^.

### Mesoscopic Modeling of Steric Interactions between Fluctuating Arms

Our model system consists of a platform of length *l*_C_+*l*_R_+*l*_L_ and two rotating arms of lengths *l*_L_ and *l*_R_ placed at distances *l*_L_ and *l*_R_ from the left and right edges of the platform, respectively, where *l*_C_ is the distance between the pivots of our arms, *l*_L_ is the length of the left arm, and *l*_R_ is the length of the right one (**Figure S8**). Both arms were treated as independent rigid bodies (they move and rotate as single entities) while the platform was held fixed. All structures were constructed from a lattice of spherical beads of diameter *σ* = 6 nm. To replicate the devices used in our experiments, we modeled the lengths of the left and right arms using 7 and 8 beads, respectively, and the left and right arm pivots were separated by a length of 9 beads. Volume exclusion was imposed between the arms and between the arms and the platform. This exclusion was modeled by means of Lennard-Jones potentials with a potential well depth *ε* = 0.001 *k*_B_*T*, a size parameter of *σ*, and a cutoff distance of *σ*, which makes the potential strictly repulsive. To keep the axes of the arms in place, bonds between the axis of the arms and the platform were introduced. Angle potentials were introduced to prevent the arms from exhibiting out-of-plane rotations.

The two arms were also subjected to separate angle-dependent potential fields *U*(*θ*) to model the hinge constraints that caused the non-uniform distribution *P*(*θ*) of hinge angles exhibited by the arms in the SteriDyn device in the free hinge condition. Specifically, we determined this field using *U*(*θ*) = −*k*_*B*_*T* ln *P*(*θ*) + *U*_0_, where *U*_0_ is a constant chosen to make *U*(*θ*;) = 0 at its minimum. These penalties were defined in LAMMPS by means of interpolation tables from potential energy and force values determined from our probability density functions. The values of *P*(*θ*) were determined by fitting experimentally-determined distributions to a Kernel distribution function defined by a normal smoothing function and a bandwidth of 1 for LF, RF, LC, RC, LC_1_, RC_1_, LC_2_, RC_2_, and to a Gaussian mixture distribution with 4 components for LC_3_, RC_3_, LC_3(HGC)_, RC_3(HGC)_B as the stiffness of these hinges leads to a narrow distribution near its maxima which was better fitted by a Gaussian function. Comparison between experimental and fitted distributions are shown in **Figure S9**. Initial positions (angles) of our arms were chosen according to a Boltzmann distribution, i.e., with a probability proportional to exp(−*U*(*θ*) /*k*_*B*_*T*), and systems where the initial conformations sterically clash were eliminated from the start.

Langevin dynamics simulations of our DNA origami devices were implemented using the Large-scale Atomic/Molecular Massively Parallel Simulator (LAMMPS) software. A Debye length of 0.3 nm was considered to model a high salt concentration of 1 M. A dielectric permittivity of 80 was used. All simulations were performed in the NVT ensemble with the temperature of the system maintained at *T* = 298 K by means of a Langevin thermostat. An integration timestep of 0.1 ps was selected, along with a damping factor of 10 ps and a mass of 5 grams/mole per bead. Simulations were run for 100 million timesteps (for a total of 10 μs), and the output was saved every 200,000 steps. The angle θ of each arm was tracked during the whole simulation time. To estimate the “true” time unit of our simulations, we fitted the mean square displacement of simulation angles to MSD(*t*) = *D*_*R*_*t*, yielding rotational diffusion coefficients *D*_*R*_ of 222.9±19.2 ns^-1^ and 222.8±21.2 ns^-1^ for the left and right arms, respectively, from 50 simulations. These coefficients are 2 million times larger than theoretical estimates of *D*_*R*_ ≈ 0.1 μs^-1^ for cylindrical structures given by *D*_*R*_ = 3*k*_*B*_*T*(ln(*L*/*d*) + *δ*)/*πη*_0_*L*^3^, where *L* = 40 nm is the length of the arms, *d* = 6 nm their diameter, *η*_0_ = 8.9×10^−4^ Pa s is the viscosity of water at room temperature, and *δ* = −0.622 + 0.917 *d*/*L* − 0.05(*d*/*L*)^2^ is a fitting function. Thus, our effective simulation time is of the order of 20 s and the simulation timestep is equivalent to 0.1 μs. Such highly coarse-grained (mesoscopic) treatment of our devices allows us to study arm-arm interactions within reasonable computational times. Comparison between our model and experiments for LC and multiple different right arms are shown in **Figure 3** and **Figure S10**.

### Derivation of Dead Zone

While engineering of hinges and overhangs provides control over the conformations sampled by the rotating arms of our devices, the separation between these arms influences their possible conformations as well. Approximating the left arm by the line segment bounded by A=(0,0,0) and B=(*l*_*L*_cos (*θ*_*L*_), *l*_*L*_sin (*θ*_*L*_), 0), and the right arm by the line segment bounded by C=(*l*_*C*_,0,0) and D=(*l*_*C*_-*l*_*R*_cos (*θ*_*R*_), *l*_*R*_sin (*θ*_*R*_), 0), with θ_L_,θ_R_ ∈ [0,π], we may implicitly define the line aligned with the left arm by those points P such that h_L_(P)=***AB***×***AP***=0, where × is the cross-product. Similarly, we may define the line aligned with the right arm by means of h_R_(P)=***CD***×***CP***=0. Both line segments intersect when h_L_(C)·h_L_(D)≤0 and h_R_(A)·h_R_(B)≤0, where · is the dot product. Replacing our expressions for the endpoints in these expressions, steric interactions between these arms prevent them from being found at angles such that

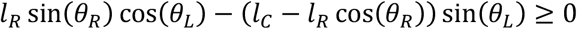

And

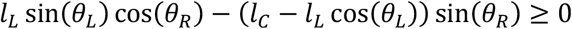

When *l*_*L*_ + *l*_*R*_ ≥ *l*_*C*_. In such case, *l*_*L*_=*l*_*R*_=0 is a trivial solution of this system of equations, as both arms would overlap when fully latched. Note that, while this solution is unique for *l*_*L*_ L *l*_*R*_ = *l*_*C*_, increasing *l*_*L*_ + *l*_*R*_ − *l*_*C*_ increases the number of solutions, i.e., the size of the conformational space both arms would intersect in.

### Mesoscopic Simulations of Steric Interactions Modulating Latching of Arms

Our previous mesoscopic simulations do an adequate job of capturing experimentally observed dynamics in systems where at least one of the arms is devoid of overhangs. However, modeling the dynamics of overhang-bearing arms by means of effective stiffer hinges is inadequate, as the prolonged residence times of the bound arms dominate the measured probability densities *P*(*θ*), obscuring the statistics of conformations outside latched states. Highly competitive systems sterically force their arms to explore less favorable states. Sampling of such states by means of poorly characterized probability densities *P*(*θ*) would then result in an inaccurate description of the dynamics of the devices. A more accurate treatment of highly competitive systems can be implemented by assigning experimentally informed binding and unbinding probabilities *p*_on_ and *p*_off_ between the arms and the platform at an angle *θ*_0_, so that the dynamics for *θ* > *θ*_0_ (“open state”) follow the energy landscape *U*(*θ*) of the corresponding arm in the absence of overhangs, while the dynamics for *θ* ≤ *θ*_0_ (“latched state”) are consolidated into a partially absorbing boundary. This treatment, which we implement in LAMMPS using the fix bond/react command, assumes overhangs have no effect on the single arm dynamics besides those associated to the binding reaction itself. As the stability of a set of overhangs is characterized by the equilibrium constant *K* = *p*_on_/*p*_off_B tuning this parameter to measurements of the percentage of latched states for independent arms would result in a better characterization of the dynamics of highly competitive systems.

In these simulations, the effects of overhangs were modeled by binding and unbinding the arms to fixed “reaction” beads placed at locations such that *θ* = *θ*_0_ when an arm is in contact with the surface of its respective bead, so that dynamics for *θ* > *θ*_0_ follow the energy landscape *U*(*θ*) of the corresponding arm in absence of overhangs, and dynamics for *θ* ≤ *θ*_0_ are consolidated in our partially absorbing boundary. Initial conformations of the arms were selected following the Boltzmann distribution of the corresponding arm in absence of overhangs limited to the interval [0B180°]. Systems where the initial conformations sterically clash were eliminated from the start. Reaction beads were modeled as spherical beads of diameter *σ* = 6 nm (see **Figure S11A**). Volume exclusion was imposed between an arm and its associated reaction bead by means of a Lennard-Jones potential with a potential well depth of *ε* = 0.001 *k*_B_*T*, a size parameter of *σ*, and a cutoff distance of *σ*. Left and right arms can only bind to their respective reaction beads, and they sterically interact with these beads in such a way that allows for binding but not for penetration. The left arm does not feel the presence of the reaction bead associated with the right arm, and the right arm does not feel the presence of the reaction bead associated with the left arm.

Binding and unbinding reactions were assessed every 100 timesteps. During each of these assessments, a random number between 0 and 1 was selected, and if this number was smaller than our desired probability (*p*_on_ for binding, *p*_off_ for unbinding), the corresponding reaction took place. A bound arm was kept fixed to the reaction bead by means of a bond of strength 0.0178 eV/Å^2^. As we are interested in thermodynamics and not kinetics, we focused on the equilibrium constant *K* = *p*_on_/*p*_off_ instead of the probabilities themselves. Hence, we considered a fixed binding probability *p*_on_ = 0.1 and fit *p*_off_ to match single arm measurements with different overhangs (see **Figure S11B**). While binding was evaluated when the arm was within a distance *σ*/2 of the center of the reaction bead, we evaluated unbinding when the arm was within a distance 3*σ*/4 to cater for the (infinitesimal) displacement of the bound arm.

Simulating the dynamics of LC|RF and LF|RC with different unbinding probabilities for the left and right arms, respectively, and defining an arm to be latched when θ≤ θ_0_L5° with a threshold of θ_0_=15°, the percentage of latched left arms for LC|RF and the percentage of latched right arms for LF|RC were well fit by three-phase exponential decay functions with K-independent parameters, 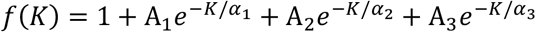. From these expressions, we obtained the equilibrium constant *K* for each set of overhangs by fitting *f*(*K*) to experimental binding percentages. These results are shown in **Figure S11C**. For the same number of overhangs, the value of *K* was found to be independent of the arm itself, as it is mostly defined by the stability of the overhang dimers. We also observe that increasing the length of the overhangs from 8 to 12 nucleotides increases the equilibrium constant *K*, although the utilized overhangs of length 12 also have a slightly higher GC content (58%) than those of length 8 (50%), which could justify the increased stability as well. Using the values of K shown in **Table S2**, 15 simulations of each device were performed (50 for strongly interacting arms) and the percentage of latched arms were compared to experiments (see **Figure 4**).

## Acknowledgements

This work is supported by the National Science Foundation primarily through Grants No. CMMI-1921881 and CMMI-1921955 and partially through CMMI-1933344. We acknowledge support from the Campus Microscopy and Imaging Facility (CMIF) at The Ohio State University for negative stain TEM imaging, and from the Texas Advanced Computing Center (TACC) at The University of Texas at Austin for providing HPC resources for carrying out the simulations presented. We also thank members of the Castro Lab, Arya Lab, and Poirier Lab for useful feedback on this work.

## Notes

### Competing Interest Statement

The authors have declared no competing interest.

