## Supporting Information for "Steric Communication between Dynamic Components on DNA nanodevices"

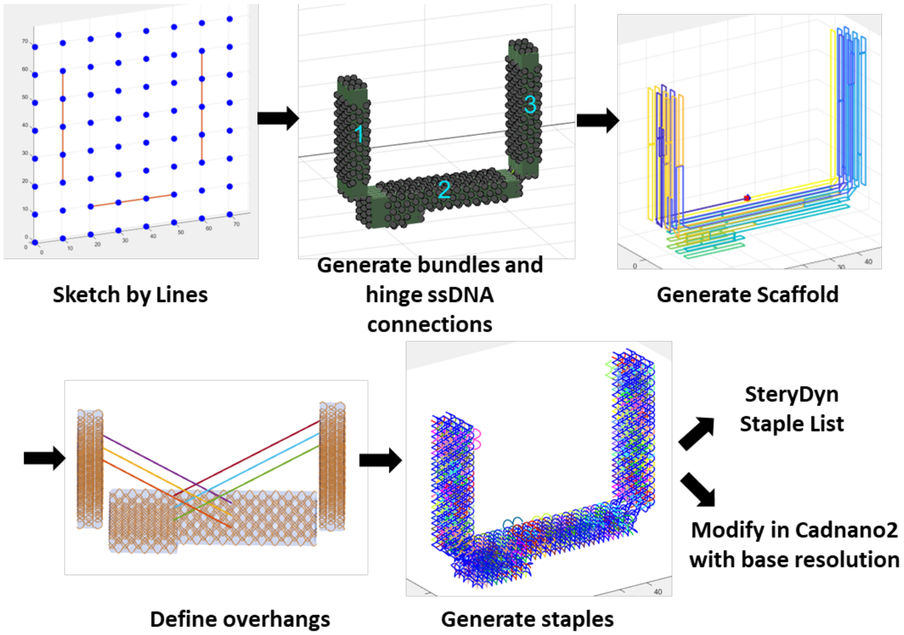

**Figure S1.** The baseline SteriDyn (free) design on MagicDNA

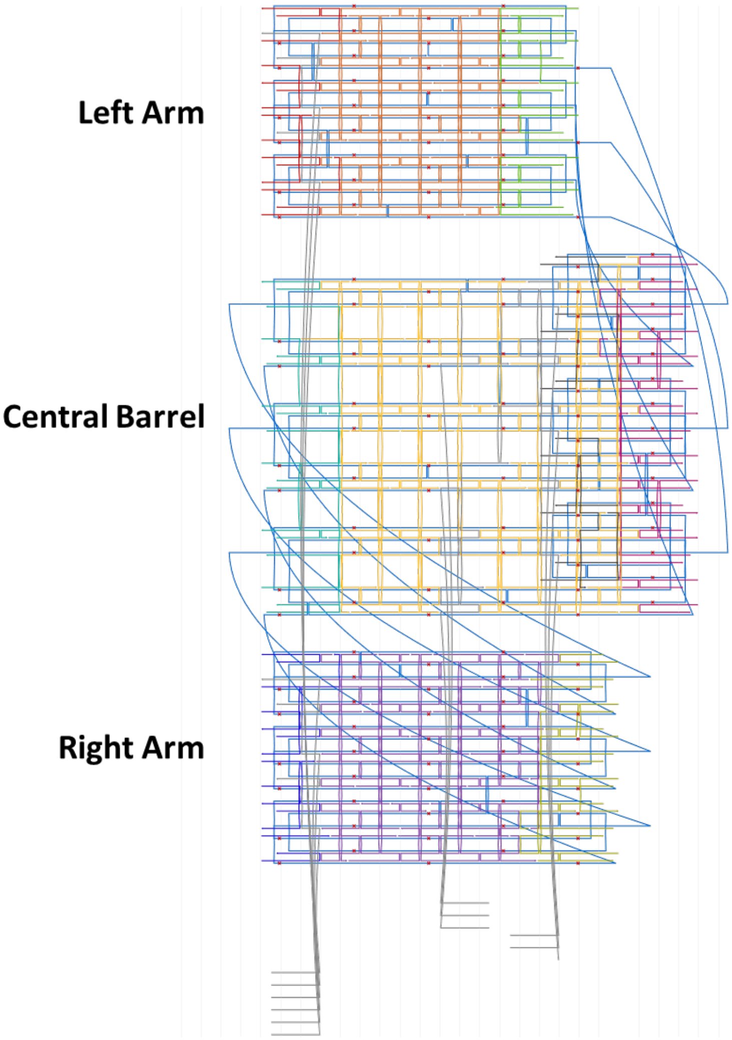

**Figure S2.** Cadnano Design Blueprint of Baseline SteriDyn (free).

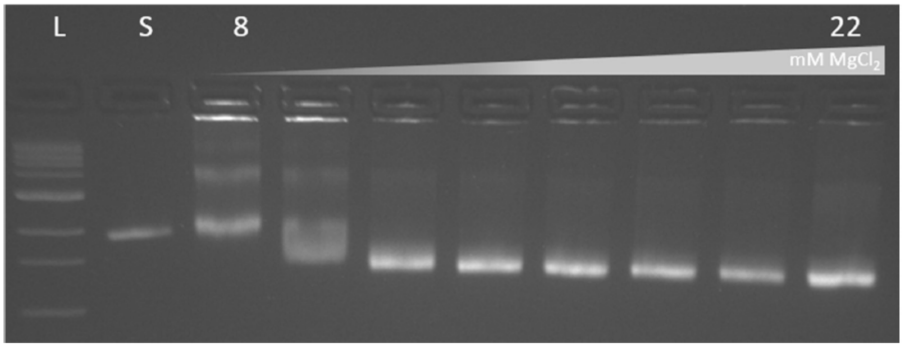

**Figure S3.** Gel electrophoresis of MgCl2 titration for baseline SteriDyn folded with [8, 10, 12, 14, 16, 18, 20, 22] mM MgCl2 concentration. L: 1kb Ladder, S: 8064 ssDNA scaffold.

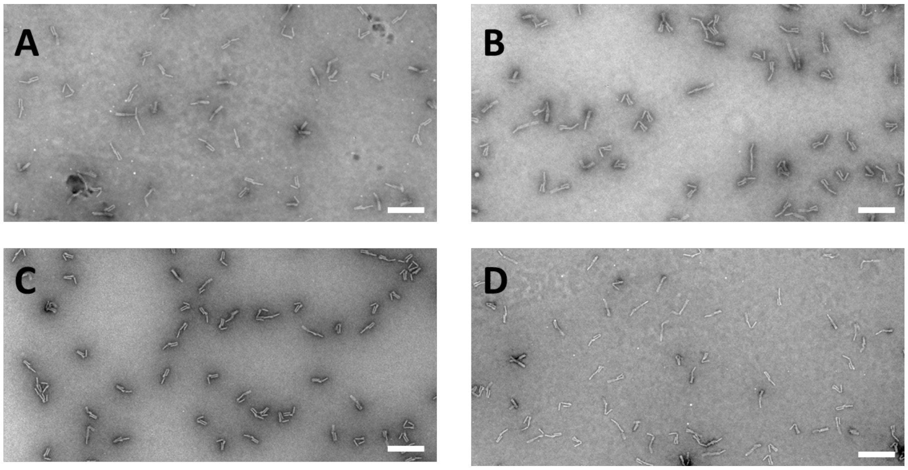

**Figure S4.** Representative TEM images of **(A)** $LC_1|RC_3$, **(B)** $LC_3|RC_1$, **(C)** $LC_1|RC_2$, **(D)** $LC_2|RC_1$. (Scale bars = 150 nm).

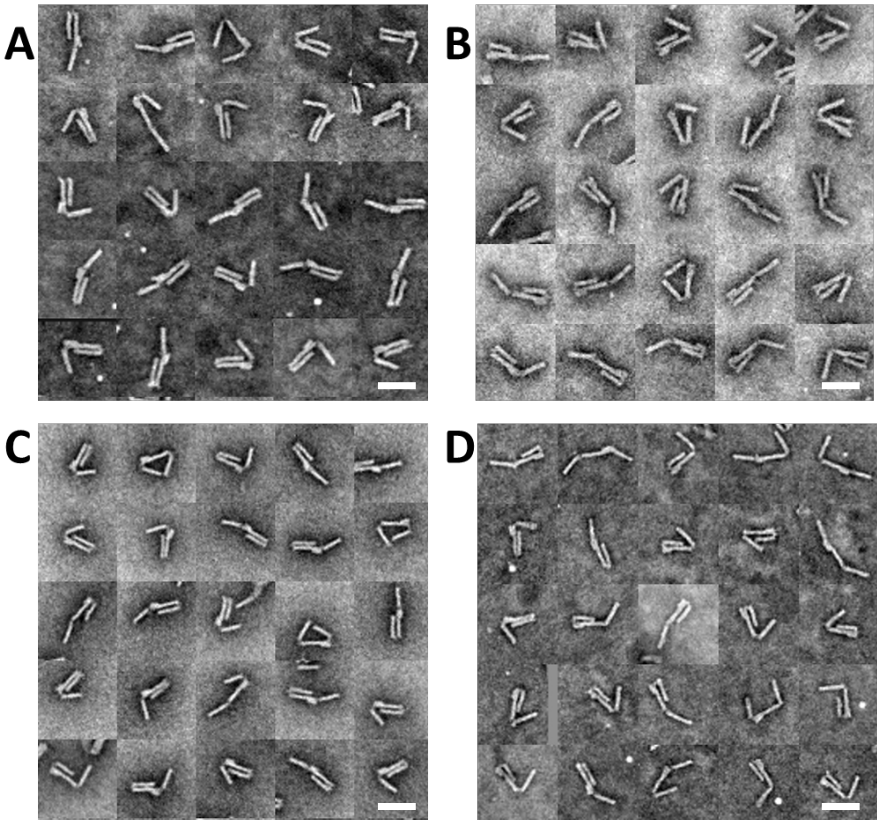

**Figure S5.** Sample TEM image galleries of **(A)** $LC_1|RC_3$, **(B)** $LC_3|RC_1$, **(C)** $LC_1|RC_2$, **(D)** $LC_2|RC_1$. (Scale bars = 50 nm).

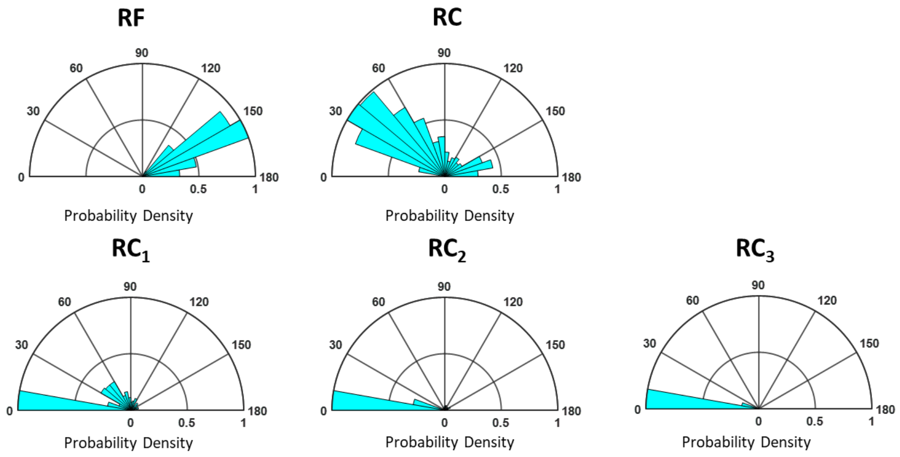

**Figure S6.** The right arm angular distribution plots with the left arm being in the free configuration. The sample size of RF, RC, RC$_1$, RC$_2$, RC$_3$ are 303, 511, 232, 185, 154.

**Commented [CC1]:** Give sample size numbers here like you do in the main figure.

**Commented [CC2R1]:** I would also plot these with the black edge lines similar to the main figure.

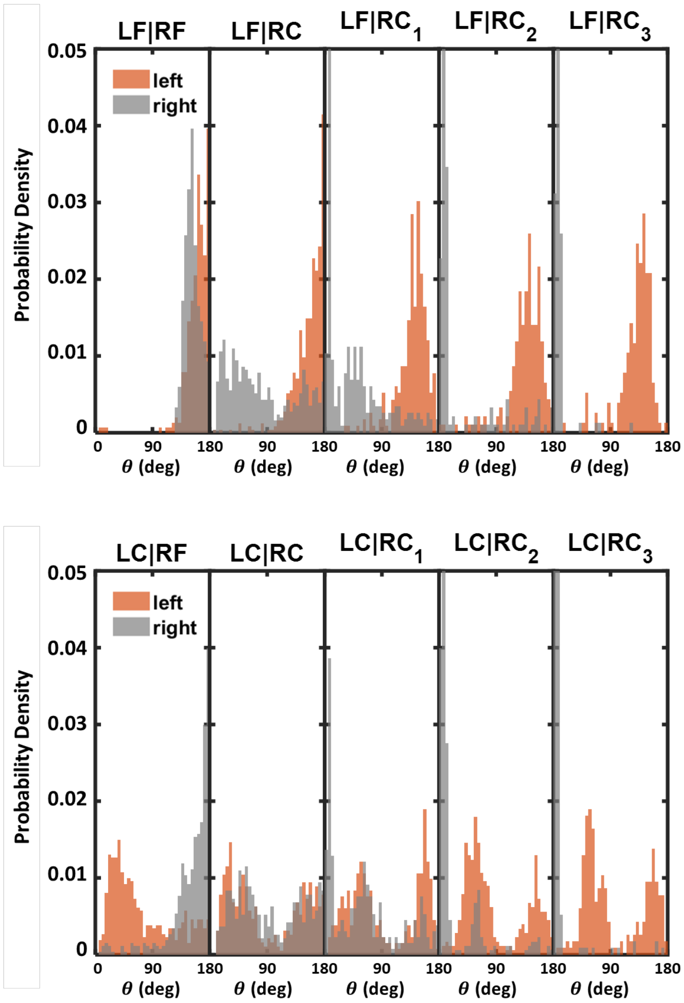
LF|RF
LF|RC
LF|RC$_1$
LF|RC$_2$
LF|RC$_3$
Probability Density
0.05
0.04
0.03
0.02
0.01
0
left
right
0
90
180
θ (deg)
LC|RF
LC|RC
LC|RC$_1$
LC|RC$_2$
LC|RC$_3$

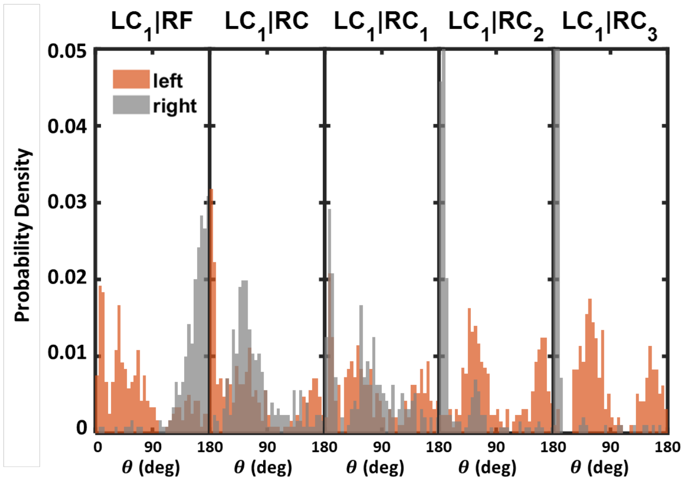
LC$_1$|RF
LC$_1$|RC
LC$_1$|RC$_1$
LC$_1$|RC$_2$
LC$_1$|RC$_3$
Probability Density
0.05
0.04
0.03
0.02
0.01
0
left
right
0
90
180
$\theta$ (deg)

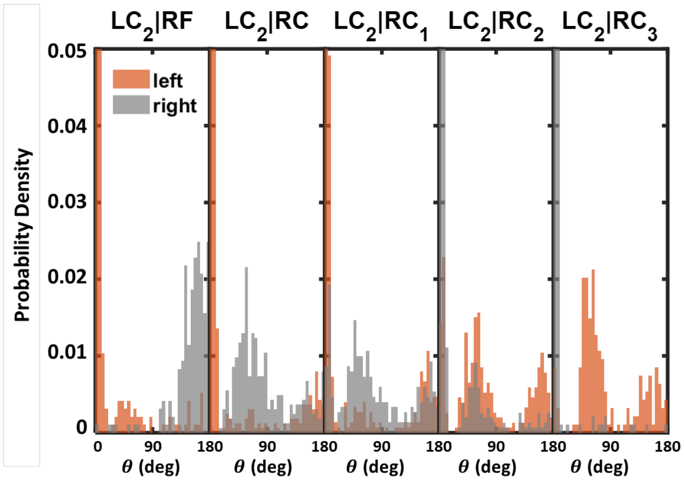
LC$_2$|RF
LC$_2$|RC
LC$_2$|RC$_1$
LC$_2$|RC$_2$
LC$_2$|RC$_3$
Probability Density
0.05
0.04
0.03
0.02
0.01
0
left
right
0
90
180
$\theta$ (deg)

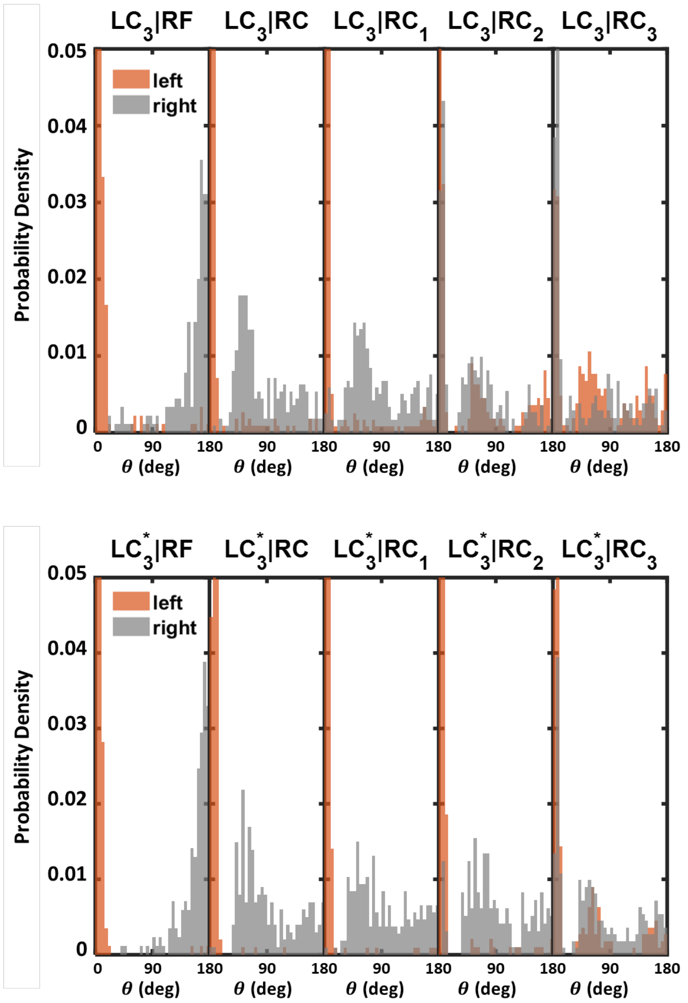

**Figure S7.** The angular distributions for all SteriDyn tested. Asterisk represents high GC conditions.

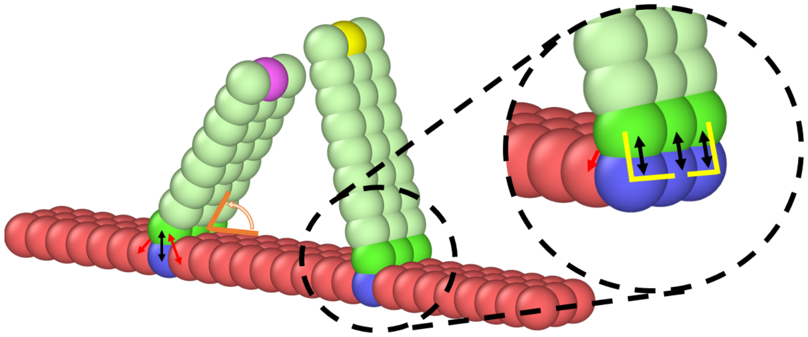

**Figure S8.** Schematic of the coarse-grained DNA origami device implemented in this work and detail of one hinge from our coarse-grained model. Arrows represent bonds that keep the base of the arms on a fixed position, and yellow lines represent the angle potentials that avoid rotation around their longitudinal axes. The stiffnesses of the bonds and angle potentials (of harmonic nature) are 0.0178 eV/A$^2$ and 10.7 eV/°$^2$, respectively, and the equilibrium lengths and angles are $\sigma$ (black), $\sqrt{2}\sigma$ (red) and 90° (yellow), respectively. The orange angle is determined between the arms, the pivot beads, and the axis of the platform. Orange angles are measured from the inside (counterclockwise for left arm, clockwise for right arm). In the rest of the manuscript, the regions of the platform placed to sterically forbid sampling of angles larger than 180° are omitted for simplicity.

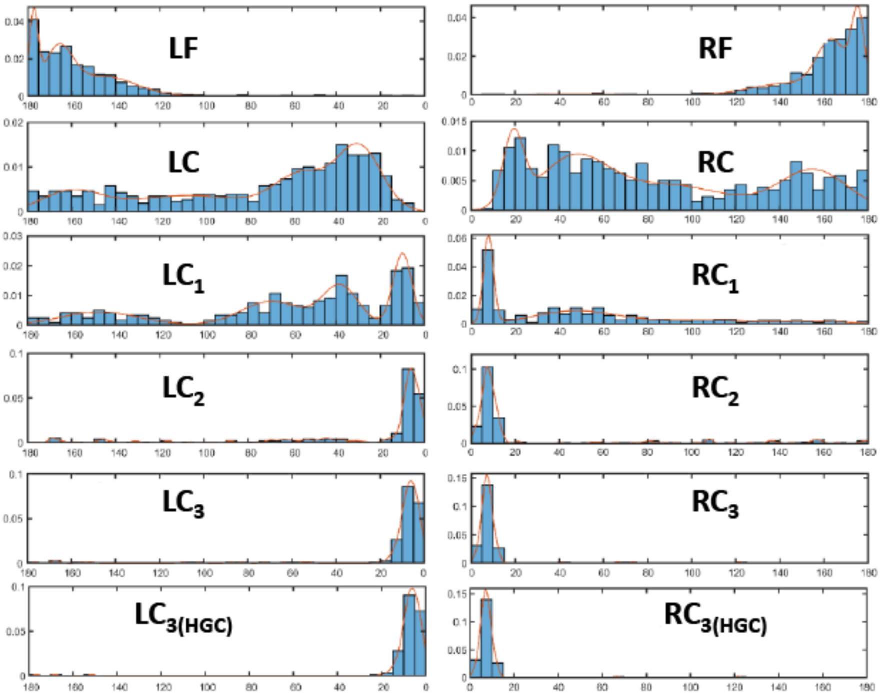

**Figure S9.** Experimental (histogram) and fitted (solid line) distributions $P(\theta)$ for the left arm of a device with a fully open right arm (RF) and for the right arm of a device with a fully open left arm (LF).

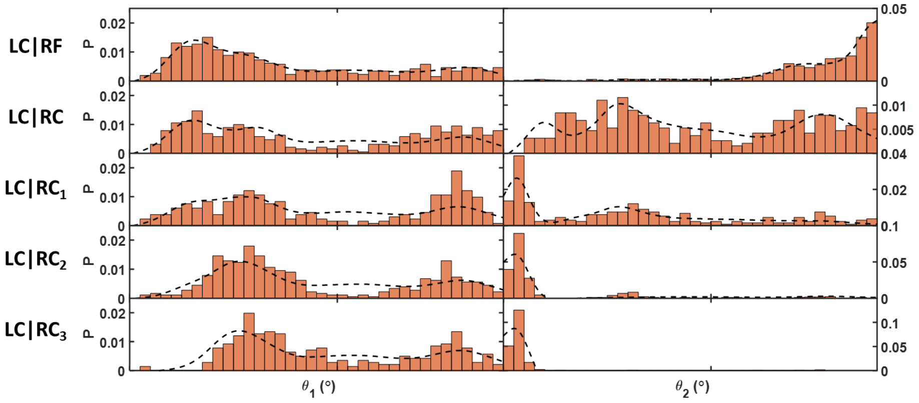

**Figure S10.** Comparison between simulated (dashed line) and experimental (bar) angular distributions for Angular phase-space of the interacting arms.

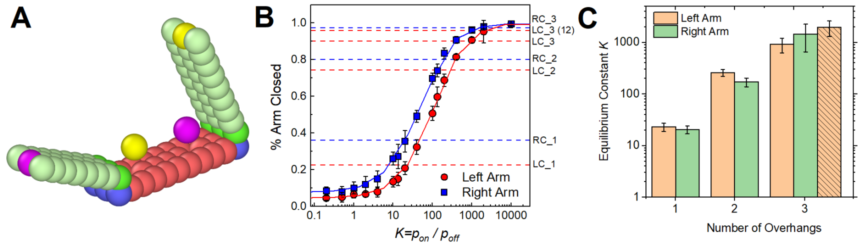

**Figure S11. (A)** Schematic of the coarse-grained DNA origami device implemented in this work, including binding sites. Note that beads of the same color (yellow, purple) are capable of binding to each other. **(B)** Simulated percentages of closed arms for different unbinding probabilities (symbols), fitting to three-phase exponential decay functions with K-independent parameters (solid lines), and experimental values of the percentage of closed arms for each arm (dashed lines). 30 simulations were run for each parameter. **(C)** Fitted equilibrium constants $K$ for arms with different numbers of overhangs (dashed bar is associated with $LC_{3(HGC)}$). Note that adding an overhang increases $K$ non-linearly (first addition by 2.5 $k_B T$, second by 1.5 $k_B T$).

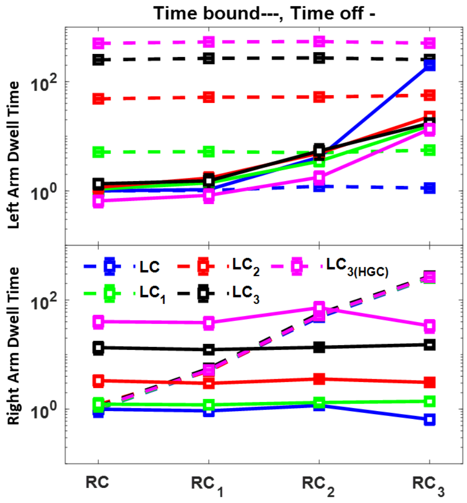

**Figure S12.** Simulation of SteriDyn dwell time for latching (bound) and unlatching(off). The dwell time are all normalized by LC|RC.

**Table S1** Sample Sizes in TEM Characterization

| Structure Name | Total Sample Size, N |
|---|---|
| LF\|RF | 303 |
| LF\|RC | 511 |
| LF\|$RC_1$ | 232 |
| LF\|$RC_2$ | 185 |
| LF\|$RC_3$ | 154 |
| LC\|RF | 521 |
| LC\|RC | 383 |
| LC\|$RC_1$ | 264 |
| LC\|$RC_2$ | 356 |
| LC\|$RC_3$ | 232 |
| $LC_1$\|RF | 240 |
| $LC_1$\|RC | 252 |
| $LC_1$\|$RC_1$ | 192 |
| $LC_1$\|$RC_2$ | 258 |
| $LC_1$\|$RC_3$ | 194 |
| $LC_2$\|RF | 193 |
| $LC_2$\|RC | 325 |
| $LC_2$\|$RC_1$ | 301 |
| $LC_2$\|$RC_2$ | 306 |
| $LC_2$\|$RC_3$ | 188 |
| $LC_3$\|RF | 180 |
| $LC_3$\|RC | 224 |
| $LC_3$\|$RC_1$ | 237 |
| $LC_3$\|$RC_2$ | 222 |
| $LC_3$\|$RC_3$ | 208 |
| $LC_{3(HGC)}$\|RF | 170 |
| $LC_{3(HGC)}$\|RC | 201 |
| $LC_{3(HGC)}$\|$RC_1$ | 213 |
| $LC_{3(HGC)}$\|$RC_2$ | 194 |
| $LC_{3(HGC)}$\|$RC_{3(HGC)}$ | 223 |
| LF\|$RC_{3(HGC)}$ | 154 |
| LC\|$RC_{3(HGC)}$ | 282 |
| $LC_1$\|$RC_{3(HGC)}$ | 131 |
| $LC_2$\|$RC_{3(HGC)}$ | 203 |

**Table S2.** Equilibrium constants used in our simulations.

| Number of Overhangs | $K$ ($p_{off}$) |
|---|---|
| 1 | 20 (0.005) |
| 2 | 200 (0.0005) |
| 3 | 1000 (0.0001) |
| 3 (HGC) | 2000 (0.00005) |

**Table S3.** Baseline SteriDyn Staple List (color coding refers to Cadnano design)

| **Staple Names** | **Sequence** |
|---|---|
| Center Barrel - Right End - 1 | TTTTTATGTTTACCAGTCGCGGTTGTGTACATCGACATTTTT |
| Center Barrel - Right End - 2 | TTTTTTAAAAAAATCGTCAGCGTGGTTTTTT |
| Center Barrel - Right End - 3 | GCTGAGGCTTGCAGGGAGTTAAACGTACAGCGCCTTTTT |
| Center Barrel - Right End - 4 | TTTTTCTTTCGCACTCAATGTAGAAC |
| Center Barrel - Right End - 5 | GCCGGATCGGCGAAAGGCCGCAGCGACAGAATCAAGTTTG CCTTTAGCGTC |
| Center Barrel - Right End - 6 | TTTTTGCTGGTCTGGGTCCAGACGACTTTTT |
| Center Barrel - Right End - 7 | GTACCTTTAATTGCTCCTTTTGTCATTGCAGGCGTTTTT |
| Center Barrel - Right End - 8 | TTTTTCGCTGAGAAGAGTGTAATTCTTCAGCAGCAACCGCA AGCCGGACTTCC |
| Center Barrel - Right End - 9 | TAGTGTCAGCGGGGATAAGAGCGACGATAAAAACCAAAAT AGCGAGAGGCT |
| Center Barrel - Right End - 10 | TTTTTGACAATAAACAACATGTTCAGCTAATAAGGTAAACA A |
| Center Barrel - Right End - 11 | AAACCAAGTACCGGATTAAGATTTTT |
| Center Barrel - Right End - 12 | ACATAGCGATAGCTTACACTCATCGAGAACAGGAAAAACG CTCATGGAAATACCTACAT |
| Center Barrel - Center Strands - 1 | GCCAACAGGCGGGGAAACAGCGCGGTTTAC |
| Center Barrel - Center Strands - 2 | GAAAGAGACGCAGAAACAGCGGAAAACGATGTGGAGCCC CCTGCGATAGTAAGCTGAATA |
| Center Barrel - Center Strands - 3 | GGCACCGCTTCAAATATTTAAAACGTGCTCGTCAGATACAT AACGCCAAAAGGAATTA |
| Center Barrel - Center Strands - 4 | AAGAGTTAAAATGTAACGCCGTTCAGCATGAGATTTTTTC |
| Center Barrel - Center Strands - 5 | CTCTTACGACAGTATCGGCCTCCAGTTTGAGCGGTCACGTT GGTGATCA |
| Center Barrel - Center Strands - 6 | CATCGATAGCAGCACCGTATAGACTTACCTCACCCCTTTAGT CTCATAACGGA |
| Center Barrel - Center Strands - 7 | CCGGAAACTCACGGAATTTCAGAGGCTGATTGTCGTCGCTG GCAGCCTTCCATG |
| Center Barrel - Center Strands - 8 | CCGTAAAAAAAGCCGCGCAGTTGGCCGGAATTTGTGAGAG AATCAGTTTTTGCG |
| Center Barrel - Center Strands - 9 | AAAGTTTCAAACTTGTGAAGGGATAGCTCGTCACCAAACAG CAT |
| Center Barrel - Center Strands - 10 | TTTTCGCACTCTGTTGGGAAGGGCGACATTTGGGAGG |
| Center Barrel - Center Strands - 11 | ATTACCCGGTTGCTGTAATAGCAAACAAAAGCAATA |

| | |
|---|---|
| Center Barrel - Center Strands - 12 | GGATCGTCACCCTCGTTGCGGTATGTGGAGGTGACG |
| Center Barrel - Center Strands - 13 | GCTTTGAGGACTACTAATGCATAAACATTGCGAACGGGC |
| Center Barrel - Center Strands - 14 | AAGTTTCCATTAATCATCAGTAATCGTTACAGACGATGAGAATCGACGCTCAAGGC |
| Center Barrel - Center Strands - 15 | TACAATAATCAGCAATTCTACATATAACTCGCAAGATACTTCTTGCGAGGCGTTTTAGCG |
| Center Barrel - Center Strands - 16 | ACGTGAATGCCATCCAGCAAATTTATCATTGCAACAAG |
| Center Barrel - Center Strands - 17 | TCGTCCCGTTCCGGCAAACGCCTGGTGCCG |
| Center Barrel - Center Strands - 18 | CCGTTGGTTTCTTACGGCCAGACGGCTACATTACCATTAGCAAGG |
| Center Barrel - Center Strands - 19 | CATCATATGTACAATTTTTGCTGTAGCCAGGGGACGTCGGA |
| Center Barrel - Center Strands - 20 | CAGAGTCATTTTTGCGGATGCGCCAGCCAAAATCATCCAAGAACCGCCTGT |
| Center Barrel - Center Strands - 21 | TATCATAACCCTCAGCAACGGATATCTCCGTGAAATTTCTGCTCATTTGCC |
| Center Barrel - Center Strands - 22 | CGAGGCGCTGGTAAATACATTTCGCAACACCTTTTTATGGTAATATCATTAC |
| Center Barrel - Center Strands - 23 | GCAGCATAAAGTACCGACAAGCAGAACGGGGTATT |
| Center Barrel - Center Strands - 24 | GTCAATAACCAGAGGCATTTTCGAGGTCCTGAAATC |
| Center Barrel - Center Strands - 25 | TTGGAGTAGATTAACTATATGAGTAGATATAGAAGGCTTATCCGG |
| Center Barrel - Center Strands - 26 | GCGAGCATATTTAACAACGCAATAAGAATAAACACCGGAATCAT |
| Center Barrel - Center Strands - 27 | CATAAAAGCCCCTGGTGTAGGGTTTTTTTCATGAATTAGAG |
| Center Barrel - Center Strands - 28 | AGAGCTTACGGCAGCCGGGTAGCACATCGATGAAGGGT |
| Center Barrel - Center Strands - 29 | TAAAGCACTTGCCTGTAAATGTAAGGCGTTAAATTTA |
| Center Barrel - Center Strands - 30 | ATTCCCAGCGCATGTAGCAACAAAGAACTTAATGGTCCTGTTTAG |
| Center Barrel - Center Strands - 31 | GAATACCGTCGGTGGTGCCACCGGCCAGCACTGTTGGCCACGGGAGCGAAAGTGAAAC |
| Center Barrel - Center Strands - 32 | TTATCAACAATAGATAACCAGTAATGAGACTACGCAACCAGCTTAATTGAGCAACAC |

| | |
|---|---|
| Center Barrel - Center Strands - 33 | CATCCTCAACATGTGGGTTATATTAGTTTGCATGTTTTAAATATGCAAC |
| Center Barrel - Center Strands - 34 | AATTACTCTTAATTGAATCCAAAGTTGATTCTGGAAGAGGAATAC |
| Center Barrel - Center Strands - 35 | AAAAGTTGAAATACCGACCGTTAAGAACTGATTAGT |
| Center Barrel - Center Strands - 36 | GGCTGTCTTTCCCGTAGGAATCCAGAACAATATTACGCTT |
| Center Barrel - Center Strands - 37 | CGAGCATCAAATCAGAAGAACTCAAACTATCTAGCTCAAACCATTAGTGGGTAAATTT |
| Center Barrel - Center Strands - 38 | CAAGCCGTTTTTATTTTCATTTATCATTAGGTCTGAAAGA |
| Center Barrel - Center Strands - 39 | TATTCGTGATAAACTGATGCAAGAATCGCCTGAAAAGGTGG |
| Center Barrel - Center Strands - 40 | AACCTCCCACCTAAATGCGAGAAATAAAGCCAATGA |
| Center Barrel - Left End (Indicator/inward) - 1 | TTCTCCGTGGGAACAAACGGCGGATTTTT |
| Center Barrel - Left End (Indicator/inward) - 2 | TTTTTATCGTAACCGTGCATCTGCAGG |
| Center Barrel - Left End (Indicator/inward) - 3 | TTTTTTTGACCGTAATGGGATATTTTTATGACCGGCAAGGCAAAGATTTTT |
| Center Barrel - Left End (Indicator/inward) - 4 | TTTTTCATCCAATAATAGATGGGCGCTTTTT |
| Center Barrel - Left End (Indicator/inward) - 5 | TTTTTATTAGCAAAATTCTAATAGTTCTACAAACAGTAGGGAGA |
| Center Barrel - Left End (Indicator/inward) - 6 | AACCGTTCTAGCTGATAAATTTTTTT |
| Center Barrel - Left End (Indicator/inward) - 7 | TTTTTTTTTGAGAGAAGTAGCATTAATTTTT |
| Center Barrel - Left End (Indicator/inward) - 8 | TATCAGTGTAGGTAAAGAGAGAGGGTAGCTATTTTTT |
| Center Barrel - Left End (Indicator/inward) - 9 | TTTTTAATGCCGTTCAAAAGGGTGA |
| Center Barrel - Left End (Indicator/outward) - 1 | TTTTTAGTAACAACCCGTAGGAACGCCATCAAAAATTTTTT |
| Center Barrel - Left End (Indicator/outward) - 2 | TTTTTTTTTTTAACCAACGCTATTACGCCAGCTGGCTTTTT |
| Center Barrel - Left End (Indicator/outward) - 3 | CATTAAAGGTGAATTATCACCGTCACCGACTTGAGCTCGGTGCGTAAGTTGGTCGC |
| Center Barrel - Left End (Indicator/outward) - 4 | TTTTTAATTCGCGTCTCTTTATTTCAATTTTT |

| | |
|---|---|
| Center Barrel - Left End (Indicator/outward) - 5 | TTTTTGAAAGGGGGATGTGCTGCAATACGTAATGCCACTACGAA |
| Center Barrel - Left End (Indicator/outward) - 6 | TTTTTAATCGGTTGTACGGAGAAGCGGCCTTCTTAAATCAGCTCATTTTT |
| Center Barrel - Left End (Indicator/outward) - 7 | TTTTTAGCATGTCAATCAGATGCCGGGTTACCTGCATTTTT |
| Center Barrel - Left End (Indicator/outward) - 8 | GGAACAACATTATTACAGGTAGAAAGATACGGGTAAAAGGCGATGGC |
| Center Barrel - Left End (Indicator/outward) - 9 | TTTTTCGCAAGGATAAAAATGAGTAATCGTAAAACTTTTTT |
| Center Barrel - Left End (Indicator/outward) - 10 | ACGGAGTCTGGACTTTTGCGCAAAAACACATCAACATTAAATGTGAGCGTTTTT |
| Center Barrel - Left End (Indicator/outward) - 11 | TTTTTGCCAGCGGTGCAAATATATTTTTTTTT |
| Center Barrel - Left End (Indicator/outward) - 12 | TTTTTTCACCATCAATACCTTTTTAGAACCCTCATATTTTT |
| Center Barrel - Left End (Indicator/outward) - 13 | TTTTTTTCTTACCAGTAACTTTTTCCGGTGCCGCGCGCCTGTGCACTCTGTGG |
| Center Barrel - Left End (Indicator/outward) - 14 | GAGTCTGTCCATCACGCAAATTAACCGTGTGTCACTCCCTGCATACGG |
| Center Barrel - Left End (Indicator/outward) - 15 | GAAAGGCCGGAGACAGTCAAATTTTT |
| Center Barrel - Left End (Indicator/outward) - 16 | TTTTTTATTTTAAATGCAATGCCTGCGTTATACAAATTTTT |
| Center Barrel - Left End (Indicator/outward) - 17 | TATCATATGAGTAATGGTCATTGTGATATTCAAGCCTCAGAGCATAAAGCTATTTTT |
| Center Barrel - Left End (Indicator/outward) - 18 | TTTTTAGTTAATTTCATCTTCTGGACTTGCGGGAGGTTTTGAAG |
| Right Barrel - End Strand - 1 | TTTTTGCTTTTGATGATAGTCAGTGCCTTGAGTAACATTTTT |
| Right Barrel - End Strand - 2 | TTTTTGTGCCCGTATTCAGGAGGTTTTTTTT |
| Right Barrel - End Strand - 3 | TTTTTAAGGCTCCAACTTTACCCTGATTTTT |
| Right Barrel - End Strand - 4 | TTTTTTCATATTCCTGATCACCGTACAAACAGTTAATGCCCCTTA |
| Right Barrel - End Strand - 5 | TTTTTAGTACCGCCAGCATCACCTTGTTTTT |
| Right Barrel - End Strand - 6 | TTTTTCTATTATAGTTCAAGAAAACATTTTT |
| Right Barrel - End Strand - 7 | TTTTTATTAAAAATACCGAATCTAAACCCTCA |
| Right Barrel - End Strand - 8 | TTTTTCTGAACCTCAAATATCAAACCC |
| Right Barrel - End Strand - 9 | TTTTTAAATTAATTACATTTAACCTAAAACATCGCCTTTTT |
| Right Barrel - Center Strand - 1 | ACGGGCAGGAGTGTACTGGTATAATAATTATCGGTTT |
| Right Barrel - Center Strand - 2 | GGAGTGAATAAGTTCTGCCTATTTTACCGGATGGCAATGACCATTGCATC |
| Right Barrel - Center Strand - 3 | CAACGAGCCGCCACCCTCAGGAACCGCTTCT |

| | |
|---|---|
| Right Barrel - Center Strand - 4 | AACAATGAAAGTTTAAAGCCTGGATTATCAAATGCACTTCA AATA |
| Right Barrel - Center Strand - 5 | CGTTAGTAAAGAAGGATGCCTTGAGAACCTACGGAATCGA AAGCGAACCAGACCGTCGC |
| Right Barrel - Center Strand - 6 | ACCGGAACCAGAGCCACCACCGAACCGCCAATCCTCAATTA AGAGGGCG |
| Right Barrel - Center Strand - 7 | AAACGACAATGACAACAACAATACTGCCATATCAAACGT |
| Right Barrel - Center Strand - 8 | TCATGCCGCCACGACGATTGTAGAGACA |
| Right Barrel - Center Strand - 9 | CCTCAAGATGAATTTAGCCACCACCCTCAGAAATC |
| Right Barrel - Center Strand - 10 | TGAATTCGGAACAGTATAGCCCGGAATCCTCAGAAGAAAC CAACCAGCAGAAAAGAATTG |
| Right Barrel - Center Strand - 11 | ATCAGCTAACGAGAATTCATCAAGGAACAAACCGCCACCCT GAGAGC |
| Right Barrel - Center Strand - 12 | TAAACAGCTTGAATCCCCCTACTTCTGTTTTAAAACAGGGAT AGACAACTCAACA |
| Right Barrel - Center Strand - 13 | CGCCCGCAGGTCACAGAACCAGTCGTCTTTCCAGA |
| Right Barrel - Center Strand - 14 | GATAAGTGCCGTTCATTTTGTTTGAGTCAG |
| Right Barrel - Center Strand - 15 | ATTCCACGATTAGCGGGCCCATGTATGAATATACAGTAACC GAGCTTCTCATAAATATTC |
| Right Barrel - Center Strand - 16 | TGTTAGAATGGAAGGTGAATCTCCCTCACTAAAGGAATTGC GAAGAATAGAATCTC |
| Right Barrel - Center Strand - 17 | ATTGATACCGATAACAAATAACCCTCAGTCTGTATGGGATT TTGCTA |
| Right Barrel - Center Strand - 18 | GAACCGCCACAGGTGTATTATCAGATTTCCAGTATTTAATT GTTTTTCAC |
| Right Barrel - Center Strand - 19 | CAGACCGTAACAATTTAGAAGTATTAGCTTT |
| Right Barrel - Center Strand - 20 | AAAAAGATTAACCTGAGCAAGATAAACTTTAATGTGGTCAG TTGGCAAA |
| Right Barrel - Center Strand - 21 | AAAGTTTAAACAGTTCAGAATGCTTTCGAAGCGCAGAGGA ACAAAGTTTCAGC |
| Right Barrel - Center Strand - 22 | TCGCGAATCGCGCGTATTAAAGGCACAGAATTGAGGAAGG TTA |
| Right Barrel - Center Strand - 23 | TAATTGTTATTAAAATAATGGCAGTACCAGCTGAGACT |
| Right Barrel - Center Strand - 24 | CAGCAGCTCAATATCCGCGAACTGATAGCCAATTTCATGAT GATGA |
| Right Barrel - Center Strand - 25 | CACGCTCAGAGCCACCACCCCGAGAGGGTGAT |
| Right Barrel - Center Strand - 26 | CCGCGAAAGGACAATATTTTTGAATGGCAGTACATATTATT CA |
| Right Barrel - Center Strand - 27 | ATTCGCAAGCCCAATAGGAAGTTTTGCTAAGGGTTATATTC ACAAGTTGCGCATC |
| Right Barrel - Center Strand - 28 | TATTAGAGGCGAAAATCAATATATGTGAGTGAATAGAATA CGTTCCTTTGCAACA |

| | |
|---|---|
| Right Barrel - Center Strand - 29 | TTTCAATTAGAGGAAGCATTTTGCTATAATCCTTGATATACT ATTATTCTGAAACACTTT |
| Right Barrel - Center Strand - 30 | TCAAAAATGAAAAACGAACCCCAGAAGGAAGCGGATAAAT CAAA |
| Right Barrel - Center Strand - 31 | TCAACAGTTCTGCAACAGAGGCGGTAACATTATCCCG |
| Right Barrel - Center Strand - 32 | TCTAAAATATACTTTACACCGAACAGTACCTTTTTTGCTTTTC TGTAAATCGTCGC |
| Right Barrel - Center Strand - 33 | AGGAGACCTGAAAGCGTAAACCTTGCTGAATACCAAGTTA CAATTT |
| Right Barrel - Center Strand - 34 | AATTACCTTTTTTAATGGAAACTATTAGTACAGAGGTGTGC |
| Right Barrel - Hinge End - 1 | TTTTTGATCTAAAGTTTTCCACCAGAGCCGCCGCCATTTTT |
| Right Barrel - Hinge End - 2 | GGTCATAGCCCCCTTATTAGCGTTTGCCATCTTT |
| Right Barrel - Hinge End - 3 | GCCCTCATAGTTAGCGTAACTTTTT |
| Right Barrel - Hinge End - 4 | TTTTTGCATTGACAGGAAACAGAAATATTTTT |
| Right Barrel - Hinge End - 5 | TTTTTAAACTACAACGCCGCACGTAAGGTTGAGACGCATAA CCGATATATTCGG |
| Right Barrel - Hinge End - 6 | ATAGTAAAATGTTTAGACTGGATAGCGTCCCAT |
| Right Barrel - Hinge End - 7 | TAGATTTTCTCCAACAGGTCAGGATTAGA |
| Right Barrel - Hinge End - 8 | TTTTTAAGAAATTGCGCCAGTAATAAATTTTT |
| Right Barrel - Hinge End - 9 | TTTTTCAATAGATAATACCACACGA |
| Right Barrel - Hinge End - 10 | CTGAACATCGGGACCCTTCTGCACTAACAACTAATAGATTA GAGCCGTTTTTT |
| Right Barrel - Hinge End - 11 | TTACATTGGCAGATTCACCAGTTAACGGATGAAGCAAACAG GTTTAAATTATTTTGTAGC |
| Right Barrel - Hinge End - 12 | TTTTTAGGGACATTCTGGCCAACAATTTTCCCTTAGAATCCT TGA |
| Right Barrel - Hinge End - 13 | TATTAATTAGAGATAGAAGAAACAAATTTGAGGCTGAGTTT CGTCACCAGTACTTTTT |
| Left Barrel - End Strand - 1 | TTTTTCCGGCGAACGTGGCCACAAGAATTGAGTTAAGTTTT T |
| Left Barrel - End Strand - 2 | TTTTTCCCAATAATAAGAGCTAAAGAACGTGGAGATAACCG AGAAAGGAA |
| Left Barrel - End Strand - 3 | TTTTTGTTGCTTTGAGCCCTGACGAGTTTTT |
| Left Barrel - End Strand - 4 | TTTTTCAGGCGCATAGGCTATAAGAAACAATGAAATATTTT T |
| Left Barrel - End Strand - 5 | TTTTTGCAATAGCTAGGCCAACGCGCTTTTT |
| Left Barrel - End Strand - 6 | TTTTTAAACACCAGACCTGATAAATTTTTTT |
| Left Barrel - End Strand - 7 | TTTTTTTTGAAAGAGGACAATGAATCTCTTA |
| Left Barrel - End Strand - 8 | TTTTTGGGGAGAGGCGGTTTGCGTATTGGGCGCT |
| Left Barrel - End Strand - 9 | TTTTTGTGTCGAAATCCGCGACCCGAACTGACCAACTTTTT |
| Left Barrel - Hinge End - 1 | TTTTTATAACATAAAAACAGGGAAGCGCATTATG |
| Left Barrel - Hinge End - 2 | GTCTTTTGCACGAACGGTACGCCAGAATCCTGAGAAGTGTT |
| Left Barrel - Hinge End - 3 | TTAAATCAAGATTAGTTGCTATTTCCAGAGCCTAATTTTT |

| | |
|---|---|
| Left Barrel - Hinge End - 4 | AAAATAAATAAATCCTCACACGTTTTCGCTCATTATACCAGTCAGGACGTTGGGAAG |
| Left Barrel - Hinge End - 5 | TTTTTTTTGCCAGTTACAAGCAGCCT |
| Left Barrel - Hinge End - 6 | TTTTTTTTGATGGTGGTTACAGAGAGATTTTT |
| Left Barrel - Hinge End - 7 | CTGCGGCCAGAATGCGGCGGGCGTTGAGGATCCCCTTTTT |
| Left Barrel - Hinge End - 8 | TTTTTGGGTACCGAGCTCTACGCAGGCGAAAATCCTGTTTTT |
| Left Barrel - Hinge End - 9 | TTTTTACGCTGGTTTGCCCCACCAAAAGAACCCCTGAGAAAGGTGGCA |
| Left Barrel - Hinge End - 10 | TATGTCAAAAGAACAAAAGGGCGACATTCAACCGATTGAGGG |
| Left Barrel - Hinge End - 11 | CACCAACCTAAAACGAAAGAGGTAGCAAACGTAGATTTTT |
| Left Barrel - Hinge End - 12 | TTTTTAAATACATACATAGAGTTGCAGCAAGCGGTCCTTTTT |
| Left Barrel - Center Strand - 1 | GGGAAGCGCTTAGCTTTCCTCGTTAGAAAACAAAGCAGAGTAATATTCCACAAAGT |
| Left Barrel - Center Strand - 2 | AGCGGGTAATTGAGCGCTAATATCAGAGACTCCAACCCCCGACTTCATCATGCTCATGCT |
| Left Barrel - Center Strand - 3 | AGGGCGCTGGCAAGGAACCCAATCCAAATAAGCTATGCGTG |
| Left Barrel - Center Strand - 4 | ACCAGAATTAACTGAACACCCTGAACAATCAAGAAACGATTTTAAAGAATAGCC |
| Left Barrel - Center Strand - 5 | CAAAGTCCAGTTTGGAACAATTTAAGAACAACATACGAGCTGATAAATTGGTCAGTGAA |
| Left Barrel - Center Strand - 6 | AATCGGAGCTAACTGCGCGTAACCACCACACCCGCCGAAAGCGAAAAGGGAGCGT |
| Left Barrel - Center Strand - 7 | CCGATTATTTCTGCCAGCACTCATGTGAATTA |
| Left Barrel - Center Strand - 8 | ACAGCCAGCTACAATTTTATCCTTGTAGCGGTCACGACAGGAGG |
| Left Barrel - Center Strand - 9 | CGAGATTACCAGGTGTGAAAGTGAGCTCCCCCAGCAGAAAATTCATATGGTT |
| Left Barrel - Center Strand - 10 | TTCCGAAATCGGCAATAACGGAAGAATTCGCAG |
| Left Barrel - Center Strand - 11 | CCTGTTCTGTTTCCTAAGGAAACGCGTTGCGCTCA |
| Left Barrel - Center Strand - 12 | GTCATACCGGGGGAAGGGATTTTATTTATTCTT |
| Left Barrel - Center Strand - 13 | CCGAAGCCCTTGAGTCCACTGGCTGACTTTAGAGCTATAACGTATGCGCCG |
| Left Barrel - Center Strand - 14 | AAGCAAACCTGTCGTGCCAGCCAGGGTCAATCATAAGGGAACTGCTCCATGAGATTTG |
| Left Barrel - Center Strand - 15 | TAGCTTCGCGTCATAAATCATTGTTTAACGTCAAAAAGACGGGAACGCTAACGAGC |
| Left Barrel - Center Strand - 16 | TGAGATGGTTTAAAAACAAAGTGTAAAGCCAGACGGTGGTTTT |
| Left Barrel - Center Strand - 17 | CAACTCGCTCACACTTGACAAAGTGTTGTGGCGAAAAACCGTCTAAGTCAGAGGGCGCT |

| | |
|---|---|
| Left Barrel - Center Strand - 18 | TCATTTACCCAAATCAACGTTCAGAGCGGGAACCCTAAGG |
| Left Barrel - Center Strand - 19 | CCTTATATCTTTGAAACTCACACGCAAAGAAGCTGATTGCCCTTCACCGCCTGGTGG |
| Left Barrel - Center Strand - 20 | GCATTAGACGGAAGCATAAAGTACAACGGTTACTT |
| Left Barrel - Center Strand - 21 | CTGCGGGCAACCACCACGGAATAAGTTTATTTGCGCTGG |
| Left Barrel - Center Strand - 22 | CATGATTAAGATTCGAGGAAACGCAATAAAATCCCTTCGTGAGCCCAGCCATATTAG |
| Left Barrel - Center Strand - 23 | GGTGCAGTCGGGAGATAGCCGAACAAAGTAGGGTTGGAACCGGAACTA |
| Left Barrel - Center Strand - 24 | TCTTTTCACCAGTGAGACCCGCTTTCCCTAATGATTGTTATCTTAA |
| Left Barrel - Center Strand - 25 | ACATATCGCCAAAGATACACTAAAACACTCGCGATTTTGTCA |
| Left Barrel - Center Strand - 26 | AGCCGGAACGAGTGTCACAATCAATGATTATACCAAGCGCGTTT |
| Left Barrel - Center Strand - 27 | TACCAGAAAAGAAATTAACTCCTTATTACGTAATCATGAAGAACTGACG |

**Table S4** Strut Staples, PolyT Staples, and Pinch Staples

| Staple Names | Sequence |
|---|---|
| Length 12 - Overhang Base-B-1 | GGGTCCTACTAGCGCGCCCAATAGCAAGGTAGAAACCTTAGGTTAATTTAGGCTGTTTAGCTATATTTGGT |
| Length 12 - Overhang Base-B-2 | GGGTCCTACTAGTAATGCTGGGCCTTGCACCTCCGGCAATCAATACAAGAAAAATAATATCC |
| Length 12 - Overhang Base-B-3 | GGGTCCTACTAGCGGAACGAGGGTAGCATGCCAAGCAAACCAGGCAAAGCGCCATTCGCCTAGCACCAGAG |
| Length 12 - Overhang Base-A-1 | TGACTCGCTTCGAATAACATTACGGTGTCCCAATTCCCCTTACACAAAAACAAAACGTTAATAT |
| Length 12 - Overhang Base-A-2 | TGACTCGCTTCGCACATTCAAAAGACTTCCCAGTCATGCGCAACCAGCCAGCTTTCC |
| Length 12 - Overhang Base-A-3 | TGACTCGCTTCGCCAGCAAAATCACCAGATTCAGGCCGACGTTGTAAATTGTGGAAGATTGTATAAGTCAT |
| Length 12 - Overhang Side Left-1 | CTAGTAGGACCCCTACAGGGCGCGTACTATGTTTTT |
| Length 12 - Overhang Side Left-2 | CTAGTAGGACCCTAAGGCTTCGAGCACGTTGACGGGGAAAGTTTTT |
| Length 12 - Overhang Side Left-3 | CTAGTAGGACCCTATCATCGACGAGTAGACGGTGTACAGACTTTTT |
| Length 12 - Overhang Side Right-1 | CGAAGCGAGTCAAACAAACACAGAAGCAAGCGGAATTATCATTTTT |
| Length 12 - Overhang Side Right-2 | CGAAGCGAGTCAAATCAGGTAAGGAGCCAGCGTCATACATGTTTTT |
| Length 12 - Overhang Side Right-3 | CGAAGCGAGTCAGTTGAAAATCTCCAAAAAAATTTTT |
| Length 8 - Overhang Base-B-1 | CCTACTAGCGCGCCCAATAGCAAGGTAGAAACCTTAGGTTAATTTAGGCTGTTTAGCTATATTTGGT |
| Length 8 - Overhang Base-B-2 | CCTACTAGTAATGCTGGGCCTTGCACCTCCGGCAATCAATACAAGAAAAATAATATCC |
| Length 8 - Overhang Base-B-3 | CCTACTAGCGGAACGAGGGTAGCATGCCAAGCAAACCAGGCAAAGCGCCATTCGCCTAGCACCAGAG |
| Length 8 - Overhang Base-A-1 | TCGCTTCGAATAACATTACGGTGTCCCAATTCCCCTTACACAAAAACAAAACGTTAATAT |
| Length 8 - Overhang Base-A-2 | TCGCTTCGCACATTCAAAAGACTTCCCAGTCATGCGCAACCAGCCAGCTTTCC |
| Length 8 - Overhang Base-A-3 | TCGCTTCGCCAGCAAAATCACCAGATTCAGGCCGACGTTGTAAATTGTGGAAGATTGTATAAGTCAT |
| Length 8 - Overhang Side Left-1 | CTAGTAGGCTACAGGGCGCGTACTATGTTTTT |
| Length 8 - Overhang Side Left-2 | CTAGTAGGTAAGGCTTCGAGCACGTTGACGGGGAAAGTTTTT |

| | |
|---|---|
| Length 8 - Overhang Side Left-3 | CTAGTAGGTATCATCGACGAGTAGACGGTGTACAGACTTTTT |
| Length 8 - Overhang Side Right-1 | CGAAGCGAAACAAACACAGAAGCAAGCGGAATTATCATTTTT |
| Length 8 - Overhang Side Right-2 | CGAAGCGAAATCAGGTAAGGAGCCAGCGTCATACATGTTTTT |
| Length 8 - Overhang Side Right-3 | CGAAGCGAGTTGAAAATCTCCAAAAAAATTTTT |
| Replacement - Overhang Base-B-1 | CGCGCCCAATAGCAAGGTAGAAACCTTAGGTTAATTTAGGCTGTTTAGCTATATTTGGT |
| Replacement - Overhang Base-B-2 | TAATGCTGGGCCTTGCACCTCCGGCAATCAATACAAGAAAAATAATATCC |
| Replacement - Overhang Base-B-3 | CGGAACGAGGGTAGCATGCCAAGCAAACCAGGCAAAGCGCCATTCGCCTAGCACCAGAG |
| Replacement - Overhang Base-A-1 | AATAACATTACGGTGTCCCAATTCCCCTTACACAAAAACAAAACGTTAATAT |
| Replacement - Overhang Base-A-2 | CACATTCAAAAGACTTCCCAGTCATGCGCAACCAGCCAGCTTTCC |
| Replacement- Overhang Base-A-3 | CCAGCAAAATCACCAGATTCAGGCCGACGTTGTAAATTGTGGAAGATTGTATAAGTCAT |
| Replacement - Overhang Side Left-1 | CTACAGGGCGCGTACTATGTTTTT |
| Replacement - Overhang Side Left-2 | TAAGGCTTCGAGCACGTTGACGGGGAAAGTTTTT |
| Replacement - Overhang Side Left-3 | TATCATCGACGAGTAGACGGTGTACAGACTTTTT |
| Replacement - Overhang Side Right-1 | AACAAACACAGAAGCAAGCGGAATTATCATTTTT |
| Replacement - Overhang Side Right-2 | AATCAGGTAAGGAGCCAGCGTCATACATGTTTTT |
| Replacement - Overhang Side Right-3 | GTTGAAAATCTCCAAAAAAATTTTT |
| Poly-T - Length 10 - CR - 1 | TTTTTTTTTTATGTTTACCAGTCGCGGTTGTGTACATCGACATTTTTTTTTT |
| Poly-T - Length 10 - CR - 2 | TTTTTTTTTTTAAAAAAATCGTCAGCGTGGTTTTTTTTTTT |
| Poly-T - Length 10 - CR - 3 | GCTGAGGCTTGCAGGGAGTTAAACGTACAGCGCCTTTTTTTTTT |
| Poly-T - Length 10 - CR - 4 | TTTTTTTTTTCTTTCGCACTCAATGTAGAAC |
| Poly-T - Length 10 - CR - 5 | TTTTTTTTTTGCTGGTCTGGGTCCAGACGACTTTTTTTTTT |
| Poly-T - Length 10 - CR - 6 | GTACCTTTAATTGCTCCTTTTGTCATTGCAGGCGTTTTTTTTTT |
| Poly-T - Length 10 - CR - 7 | TTTTTTTTTTCGCTGAGAAGAGTGTAATTCTTCAGCAGCAACCGCAAGCCGGACTTCC |
| Poly-T - Length 10 - CR - 8 | TTTTTTTTTTGACAATAAACAACATGTTCAGCTAATAAGGTAAACAA |
| Poly-T - Length 10 - CR - 9 | AAACCAAGTACCGGATTAAGATTTTTTTTTT |

| | |
|---|---|
| Poly-T - Length 10 - CL - 1 | TTTTTTTTTTAGTAACAACCCGTAGGAACGCCATCAAAAATTTTTTT TTT |
| Poly-T - Length 10 - CL - 2 | TTTTTTTTTTTTTTTTAACCAACGCTATTACGCCAGCTGGCTTTTTTTTT T |
| Poly-T - Length 10 - CL - 3 | TTTTTTTTTTAATTCGCGTCTCTTTATTTCAATTTTTTTTTT |
| Poly-T - Length 10 - CL - 4 | TTTTTTTTTTGAAAGGGGGATGTGCTGCAATACGTAATGCCACTACG AA |
| Poly-T - Length 10 - CL - 5 | TTTTTTTTTTAATCGGTTGTACGGAGAAGCGGCCTTCTTAAATCAGCT CATTTTTTTTTT |
| Poly-T - Length 10 - CL - 6 | TTTTTTTTTTAGCATGTCAATCAGATGCCGGGTTACCTGCATTTTTTTT TT |
| Poly-T - Length 10 - CL - 7 | TTTTTTTTTTCGCAAGGATAAAAATGAGTAATCGTAAAACTTTTTTTT TTT |
| Poly-T - Length 10 - CL - 8 | ACGGAGTCTGGACTTTTGCGCAAAAACACATCAACATTAAATGTGA GCGTTTTTTTTTT |
| Poly-T - Length 10 - CL - 9 | TTTTTTTTTTGCCAGCGGTGCAAATATATTTTTTTTTTTTTT |
| Poly-T - Length 10 - CL - 10 | TTTTTTTTTTTCACCATCAATACCTTTTTAGAACCCTCATATTTTTTTTT T |
| Poly-T - Length 10 - CL - 11 | TTTTTTTTTTTTCTTACCAGTAACTTTTTCCGGTGCCGCGCGCCTGTGC ACTCTGTGG |
| Poly-T - Length 10 - CL - 12 | GAAAGGCCGGAGACAGTCAAATTTTTTTTTT |
| Poly-T - Length 10 - CL - 13 | TTTTTTTTTTTATTTTAAATGCAATGCCTGCGTTATACAAATTTTTTTTT T |
| Poly-T - Length 10 - CL - 14 | TATCATATGAGTAATGGTCATTGTGATATTCAAGCCTCAGAGCATAA AGCTATTTTTTTTTT |
| Poly-T - Length 10 - CL - 15 | TTTTTTTTTTAGTTAATTTCATCTTCTGGACTTGCGGGAGGTTTTGAA G |
| Poly-T - Length 10 - Right 1 | TTTTTTTTTTGATCTAAAGTTTTCCACCAGAGCCGCCGCCATTTTTTTT TT |
| Poly-T - Length 10 - Right 2 | GCCCTCATAGTTAGCGTAACTTTTTTTTTT |
| Poly-T - Length 10 - Right 3 | TTTTTTTTTTGCATTGACAGGAAACAGAAATATTTTTTTTTT |
| Poly-T - Length 10 - Right 4 | TTTTTTTTTTAAACTACAACGCCGCACGTAAGGTTGAGACGCATAAC CGATATATTCGG |
| Poly-T - Length 10 - Right 5 | TAGATTTTCTCCAACAGGTCAGGATTAGA |
| Poly-T - Length 10 - Right 6 | TTTTTTTTTTAAGAAATTGCGCCAGTAATAAATTTTTTTTTT |
| Poly-T - Length 10 - Right 7 | TTTTTTTTTTCAATAGATAATACCACACGA |
| Poly-T - Length 10 - Right 8 | CTGAACATCGGGACCCTTCTGCACTAACAACTAATAGATTAGAGCCG TTTTTTTTTT |

| | |
|---|---|
| Poly-T - Length 10 - Right 9 | TTTTTTTTTTAGGGACATTCTGGCCAACAATTTTCCCTTAGAATCCTTGA |
| Poly-T - Length 10 - Right 10 | TATTAATTAGAGATAGAAGAAACAAATTTGAGGCTGAGTTTCGTCACCAGTACTTTTTTTTTT |
| Poly-T - Length 10 - Left 1 | TTTTTTTTTTATAACATAAAAACAGGGAAGCGCATTATG |
| Poly-T - Length 10 - Left 2 | TTAAATCAAGATTAGTTGCTATTTCCAGAGCCTAATTTTTTTTTT |
| Poly-T - Length 10 - Left 3 | TTTTTTTTTTTTTGCCAGTTACAAGCAGCCT |
| Poly-T - Length 10 - Left 4 | TTTTTTTTTTTTTGATGGTGGTTACAGAGAGATTTTTTTTTT |
| Poly-T - Length 10 - Left 5 | CTGCGGCCAGAATGCGGCGGGCGTTGAGGATCCCCTTTTTTTTTT |
| Poly-T - Length 10 - Left 6 | TTTTTTTTTTGGGTACCGAGCTCTACGCAGGCGAAAATCCTGTTTTTTTTTT |
| Poly-T - Length 10 - Left 7 | TTTTTTTTTTACGCTGGTTTGCCCCACCAAAAGAACCCCTGAGAAAGGTGGCA |
| Poly-T - Length 10 - Left 8 | CACCAACCTAAAACGAAAGAGGTAGCAAACGTAGATTTTTTTTTT |
| Poly-T - Length 10 - Left 9 | TTTTTTTTTTAAATACATACATAGAGTTGCAGCAAGCGGTCCTTTTTTTTTT |
| Pinch Staple - Length 15 - Left - 2 (inner) | AAAATAAATAAATCCTCACACGTTTTCGCTCATTA |
| Pinch Staple - Length 15 - CL - 3 (inner) | CGTCACCGACTTGAGCTCGGTGCGTAAGTTGGTCGC |
| Pinch Staple - Length 15 - CR -3 (inner) | GCCGGATCGGCGAAAGGCCGCAGCGACAG |
| Pinch Staple - Length 15 - CL- 2 (inner) | AGAAAGATACGGGTAAAAGGCGATGGC |
| Pinch Staple - Length 15 - CR - 2 (inner) | TAGTGTCAGCGGGGATAAGAGCGACGATA |
| Pinch Staple - Length 15 - CL - 1 (inner) | TTAACCGTGTGTCACTCCCTGCATACGG |
| Pinch Staple - Length 15 - CR - 1 (inner) | ACATAGCGATAGCTTACACTCATCGAGAACAGGAAAA |
| Pinch Staple - Length 15 - Right - 1 (inner) | TAACGGATGAAGCAAACAGGTTTAAATTATTTTGTAGC |
| Pinch Staple - Length 15 - left - 1 | GTCTTTTGCACGAACGGTACGCCAGAATCCTGAGAAGTGTTTTTATAATCAGTGA |
| Pinch Staple - Length 15 - Left - 2 | TACCAGTCAGGACGTTGGGAAGAAAAATCTACGTTA |
| Pinch Staple - Length 15 - Left - 3 | TATGTCAAAAGAACAAAAGGGCGACATTCAACCGATTGAGGGAGGGAAGGTAAATA |
| Pinch Staple - Length 15 - CR - 3 | AATCAAGTTTGCCTTTAGCGTCAGACTGTAGCGCGTT |
| Pinch Staple - Length 15 - CL - 3 | TTGACGGAAATTATTCATTAAAGGTGAATTATCAC |

| | |
|---|---|
| Pinch Staple - Length 15 - CR - 2 | AAAACCAAAATAGCGAGAGGCTTTTGCAAAAGAAGTT |
| Pinch Staple - Length 15 - CL - 2 | ATAAAACGAACTAACGGAACAACATTATTACAGGT |
| Pinch Staple - Length 12 - CR - 1 | ACGCTCATGGAAATACCTACATTTTGACGCTCAATCG |
| Pinch Staple - Length 15 - CL - 1 | GGCCACCGAGTAAAAGAGTCTGTCCATCACGCAAA |
| Pinch Staple - Length 15 - Right - 3 | TTCATCGGCATTTTCGGTCATAGCCCCCTTATTAGCGTTTGCCATCTT T |
| Pinch Staple - Length 15 - Right - 2 | TTGCCAGAGGGGGTAATAGTAAAATGTTTAGACTGGATAGCGTCCC AT |
| Pinch Staple - Length 15 - Right - 1 | TCTGAAATGGATTATTTACATTGGCAGATTCACCAGT |
